# Sequence-Independent In Situ Imaging of GlycoRNA in Living Cells and Tissues based on COMPASS

**DOI:** 10.64898/2026.04.01.715772

**Authors:** Bingzhi Li, Sirui Liu, Yue Chen, Shulin Wang, Xue Li, Xinyue Zhang, Ruoyu Jiao, Muhammad Sohail, Wanying Zhu, Yufan Zhang, Xing Zhang

## Abstract

Current methods for the imaging of glycosylated RNAs (glycoRNAs) focus on limited types with identified RNA sequences, making the distribution of global glycoRNAs at single-cell and tissue levels largely unexplored. To address this gap, we developed a **<u>co</u>**-**<u>m</u>**etabolic **<u>p</u>**roximity labeling **<u>ass</u>**ay (COMPASS) for sequence-independent imaging of glycoRNAs. COMPASS employs metabolic chemical reporters (MCRs) to label glycans and RNAs, followed by click-chemistry-based dye conjugation to generate Förster resonance energy transfer (FRET) signals. The MCR-enabled RNA labeling allows for sequence-independent imaging, and FRET ensures accurate localization of glycoRNA distributions. COMPASS facilitates glycoRNA imaging across multiple cell lines, cerebral organoids, and mouse tissues. We observed the downregulation of glycoRNAs in metastatic lung nodules and upregulation in original regions of brain development in cerebral organoids, suggesting their potential association with carcinogenesis and neurogenesis. COMPASS illuminates previously invisible glycoRNAs and offers a comprehensive profile of glycoRNA distribution for studying their functions.

---

Glycosylation is an essential post-translational modification of biomolecules across life forms, primarily occurring in proteins and lipids to form glycoproteins and glycolipids^1^. This modification influences structural folding, intracellular trafficking, and immune recognition by mediating the molecular interactions of glycosylated substrates^2, 3^. Recent discoveries have identified RNA as a novel substrate within the cellular glycosylation network, leading to the formation of glycosylated RNAs (glycoRNAs)^4^. These molecules are displayed on the cell surfaces and are characterized by small RNAs conjugated with N-glycans terminated in sialic acids^4^. The discovery of glycoRNAs suggests the potential existence of previously unrecognized glycosylation and RNA modification pathways, both of which hold biological and clinical implications^5, 6^. Despite these insights, the function of glycoRNAs remains poorly understood. Emerging evidence indicates that glycoRNAs mediate neutrophil–endothelial interactions and modulate immune responses, positioning them as promising therapeutic targets in inflammation and cancer ^7–10^. Further elucidation of glycoRNA functions relies on the development of techniques for analyzing their distributions and abundances.

GlycoRNAs can be detected with *in vitro* techniques, for example, a modified Northern Blot assay involves metabolic labeling, RNA gel separation, dye conjugation, and infrared gel imaging^4^. The Flynn group later introduced an improved technique termed rPAL, which replaces metabolic labeling with periodate oxidation and aldehyde labeling^11^, enhancing the sensitivity of glycoRNA detection. Additionally, by adjusting the labeling and conjugation methods, the detection of glycoRNAs can be achieved with the integration of mass spectrometry and RNA sequencing^11–13^. However, these *in vitro* techniques exhibit limitations, including the need for substantial cell numbers and the risk of glycoRNA degradation during prolonged treatments^7^. These limitations render them inadequate for examining glycoRNA distributions and elucidating their functions.

Alternatively, glycoRNAs can be detected using fluorescent imaging techniques. The Lu group pioneered glycoRNA imaging by developing ARPLA, which exploits the proximity of sialic acid aptamers and RNA-binding probes to trigger rolling circle amplification (RCA)^8^. The Ju group proposed a second-generation hierarchical coding strategy (HieCo-2), which integrates a hybridization chain reaction (HCR) for the visualization of Y5 glycoRNAs^14^. More recently, the Xu group developed a dual-recognition Förster resonance energy transfer (drFRET) platform that combines RNA-binding probes with sialic acid aptamers, enabling the imaging of small extracellular vesicle (sEV)-derived glycoRNAs^15^. Despite these advances, all reported approaches rely on DNA nanotechnologies, which require sophisticated probe design and precise control of hybridization on the cell surface. Moreover, RCA and HCR generate long DNA fibers (up to several hundred nanometers)^16–18^, which can displace the fluorescence signal from the actual site of glycoRNA, thereby compromising the accuracy of glycoRNA localization. A more fundamental limitation arises from the sequence-dependence of these assays: because they rely on complementary probes targeting the RNA moiety, their applicability is restricted to glycoRNAs with known sequences. Therefore, there is a need for the development of robust imaging techniques capable of profiling comprehensive glycoRNA information.

In this work, we developed a sequence-independent glycoRNA imaging technique, termed **<u>co</u>**-**<u>m</u>**etabolic **<u>p</u>**roximity labeling **<u>ass</u>**ay (COMPASS). COMPASS integrates co-metabolic labeling of RNAs and glycans, click chemistry-based dye conjugation, and FRET imaging. COMPASS utilizes metabolic chemical reporters (MCRs) to incorporate into the biosynthesis of RNAs and glycans, labeling them with alkyne and azide (N_3_) groups, respectively. A selection of dyes capable of generating FRET signals is then conjugated to the alkyne and N_3_ groups through click chemistry approaches. Given that FRET occurs within a 10 nm range^19^, the proximity of RNAs and glycans within a single glycoRNA molecule activates the FRET signal, facilitating the imaging of glycoRNAs. In contrast to earlier DNA probe-based approaches^14, 20^, some of which also exploited FRET^15^, COMPASS employs 5-ethynyluridine (5-EU) for RNA labeling and eliminates bulky recognition elements, thereby improving localization accuracy and ensuring sequence-independent detection. This strategy demonstrated broad applicability by enabling glycoRNA imaging across multiple cell lines, human cerebral organoids, and mouse tissues, representing the first approach capable of comprehensive profiling at both the cellular and tissue levels.

## Results

### Development of COMPASS for cell surface glycoRNA imaging

COMPASS incorporates two functional moieties: (1) MCRs including Ac_4_ManNAz and 5-EU. Ac_4_ManNAz incorporates into glycan biosynthetic pathways to label sialic acids in glycoRNAs with N_3_^21^. 5-EU incorporates into RNA *de novo* synthesis, labeling U with alkyne groups in newly generated RNAs^22^; (2) FRET pairs, consisting of energy donor N_3_-AF488 and energy acceptor DBCO-AF647.

As illustrated in **Fig. 1a**, cells were co-incubated with MCRs, allowing the glycans and RNAs to be labeled with N_3_ and alkyne groups, respectively (**Step 1**). Subsequently, DBCO-AF647 was introduced to react with N_3_ through copper-free click chemistry, thereby localizing to the cell surface sialic acid (**Step 2**). After washing to remove the free DBCO-AF647, RNAs were targeted using copper(I)-catalyzed azide-alkyne cycloaddition (CuAAC), where N_3_-AF488 was linked to the alkynes on U (**Step 3**). As AF488 and AF647 are in proximity (<10 nm), the emission from AF488 excites AF647, generating a FRET signal for in situ glycoRNA imaging. In this framework, AF488 indicates RNA localization, AF647 identifies sialic acids, and the FRET signal visualizes glycoRNAs.

**Fig. 1.**
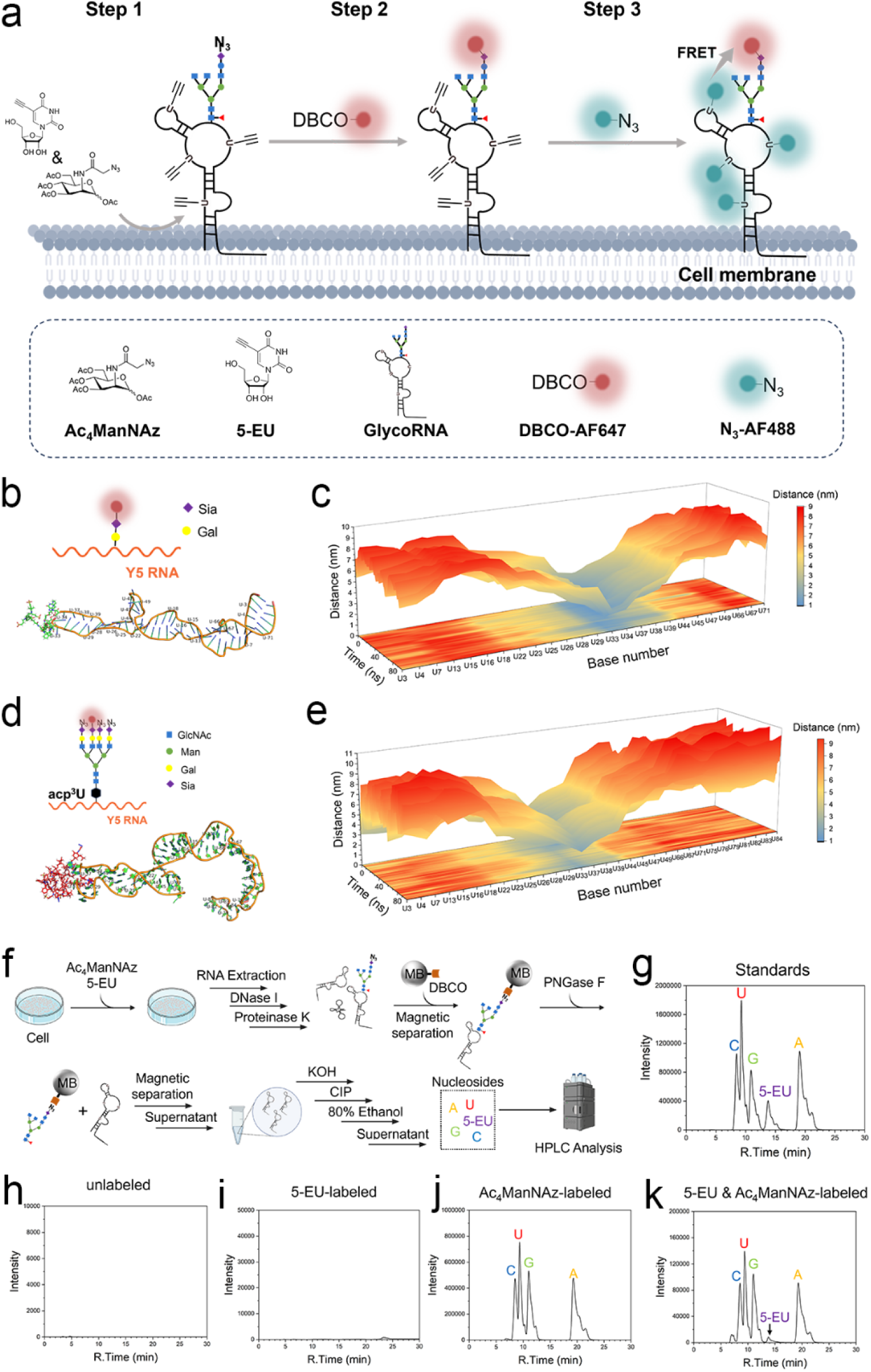
Schematic illustration of COMPASS, MD simulations, and co-metabolic labeling verifications. **a,** Functional moieties and steps of the COMPASS. **b,** A representative structure of synthetic Y5 glycoRNA modified with AF647. The Y5 glycoRNA with sialic acid terminus was labeled with N_3_ and then conjugated with DBCO-AF647 via N_3_-DBCO conjugation. This Y5 glycoRNA contains a total of 25 U bases that can be potentially labeled by 5-EU through metabolic labeling, which are marked with ordinal numbers for clarity. The chemical structure of this Y5 glycoRNA is detailed in **Supplementary Fig. 1. c,** Dynamic distances between AF647 and each U residue over a 100-ns MD simulation of the structure shown in panel **b** calculated using GROMACS. **d,** A representative structure of a physiologically relevant Y5 glycoRNA model modified with AF647. The Y5 RNA was linked to a complex N-glycan via an acp^3^U linkage, with the terminal sialic acid modified with N_3_ group and subsequently conjugated to DBCO-AF647 via N_3_-DBCO conjugation. This Y5 glycoRNA contains a total of 31 U residues, each annotated with an ordinal number for reference. The chemical structure of this Y5 glycoRNA is detailed in **Supplementary Fig. 2. e,** Dynamic distances between AF647 and each U residue over a 100-ns MD simulation of the structure shown in panel **d** calculated using GROMACS. **f,** Schematic illustration of glycoRNA enrichment using DBCO-functionalized magnetic beads (MBs), followed by on-bead RNA release via PNGase F treatment, in-solution RNA hydrolysis, and HPLC analysis. **g-k,** HPLC chromatograms of an equimolar mixture of nucleoside standards (**g**) and glycoRNA isolated from unlabeled cells (**h**), labeled with 5-EU (**i**), labeled with Ac_4_ManNAz (**j**), or co-labeled with 5-EU and Ac_4_ManNAz (**k**).

Using the structure of a synthetic Y5 glycoRNA^13^, we performed molecular dynamics (MD) simulations and found that the distance between the AF647 (attached to the glycan) and all the 25 U fluctuated within 10 nm throughout the 100 ns simulation (**Figs. 1b, 1c, and Supplementary Fig. 1**). We next extended these simulations to a more physiologically relevant model comprising a full-length N-glycan linked to RNA via 3-(3-amino-3-carboxypropyl) uridine (acp^3^U) (**Fig. 1d and Supplementary Fig. 2**)^11, 23^. The glycan structure was chosen to reflect the most abundant natural configuration identified in a recent mass spectrometry study^23^. During a 100 ns MD simulation, all U residues remained within 10 nm of AF647 for the majority of the time. Only 14 U transiently exceeded this distance, with a maximum of 11.012 nm, and such deviations accounted for less than 3.9% of the simulation period (**Fig. 1e**). These results indicate that in glycoRNA, the glycan-attached AF647 and most U have FRET-compatible proximity, supporting the feasibility of COMPASS for FRET imaging.

CCK-8 assay demonstrated that the co-treatment of MCF-7 cells with 50 μM Ac_4_ManNAz and 250 μM 5-EU for 48 h, as well as with lower concentrations, maintained cell viability above 95% (**Supplementary Fig. 3**). While increasing the concentrations to 100 μM Ac_4_ManNAz and 500 μM 5-EU significantly reduced cell viability to 86%. Therefore, based on the cytotoxicity and previous reports^24, 25^, we selected 50 μM Ac_4_ManNAz and 250 μM 5-EU for 48 h for initial COMPASS labeling.

We then investigated the direct incorporation of Ac_4_ManNAz and 5-EU into cellular glycans and RNAs, respectively (**Supplementary Figs. 4 and 5**). In MCF-7 cells treated with 50 μM Ac_4_ManNAz, sialic acids were isolated, hydrolyzed, and purified. Then these cell-derived sialic acids and their derivative (e.g., N_3_-labeled sialic acid, Neu5Az) were processed with 1,2-diamino-4,5-methylenedioxybenzene (DMB) derivatization and high-performance liquid chromatography (HPLC) analysis^26^. Cells treated with Ac_4_ManNAz exhibited a distinct peak corresponding to DMB-Neu5Az, confirming incorporation of N_3_ into cellular sialic acids (**Supplementary Figs. 4a–4d**). Meanwhile, RNAs isolated from MCF-7 cells treated with 250 μM 5-EU were hydrolyzed and analyzed by HPLC^22^, revealing peaks corresponding to 5-EU (**Supplementary Figs. 5a–5d**), confirming the effective 5-EU labeling of cellular RNA. The incorporation rates (IRs) of Ac_4_ManNAz into cellular sialic acids and of 5-EU into U were determined to be 59.02% and 3.76%, respectively, comparable to previous reports^22, 27^. Importantly, when performing co-metabolic labeling, the IRs of both MCRs remained comparable to those observed during individual labeling (**Supplementary Figs. 4e and 5e**), indicating minimal metabolic interference between these two MCRs.

The presence of glycoRNAs in MCF-7 cells was first confirmed by RNA blotting **(Supplementary Fig. 6)**^4^. To further validate the co-metabolic incorporation of 5-EU and Ac_4_ManNAz into the same glycoRNA molecules, glycoRNAs from MCF-7 cells were enriched using DBCO-functionalized magnetic beads (MBs), enzymatically treated with PNGase F to release the RNA moiety, and subsequently hydrolyzed to nucleosides for HPLC analysis (**Fig. 1f**). The 5-EU signal was identified by comparison with authentic standards, providing direct evidence that both 5-EU and Ac_4_ManNAz are co-metabolically incorporated into glycoRNAs at the cellular level (**Figs. 1g–k**). Notably, the IR of 5-EU in glycoRNAs reached approximately 10.39%, markedly higher than that in total RNA (3.76%), indicating preferential incorporation of 5-EU into glycoRNA-associated transcripts. GlycoRNAs are primarily composed of small noncoding RNAs^28, 29^, which are predominantly transcribed by RNA polymerases II and III^30, 31^. Previous studies have shown that 5-EU is more efficiently incorporated into transcripts generated by RNA polymerases II and III^22^, which explains the elevated 5-EU incorporation in glycoRNAs.

We then utilized confocal laser-scanning microscopy (CLSM) for the in-situ imaging of glycoRNA based on COMPASS (**Fig. 2a)**. With complete FRET pairs introduced, only the AF647 signal corresponding to sialic acid was detected without 5-EU addition (**Fig. 2a**, Row 1), and similarly, only the AF488 signal representing RNAs was observed without Ac_4_ManNAz addition (**Fig. 2a**, Row 2). In groups that underwent co-metabolic labeling and were treated with either DBCO-AF647 or N_3_-AF488 individually, the signals were observed based on the specific dye present; however, no FRET signal was detected under these conditions (**Fig. 2a**, Rows 3 and 4). Under full COMPASS operation, we observed an intensified FRET signal on the membranes of all MCF-7 cells (**Fig. 2a**, Row 5).

**Fig. 2.**
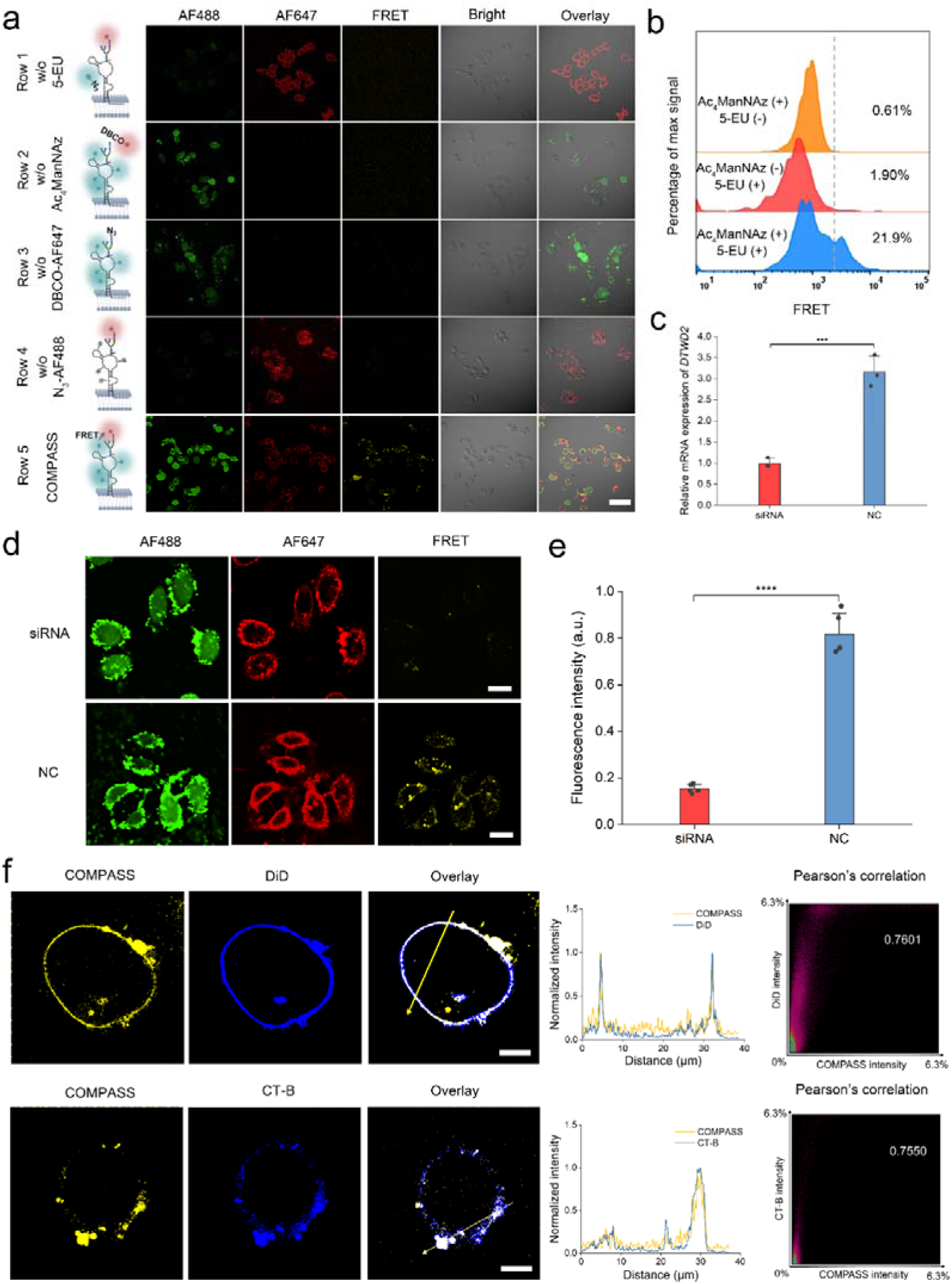
Feasibility of COMPASS for glycoRNA imaging in MCF-7 cells. **a,** CLSM images of glycoRNAs on the surface of MCF-7 cells with COMPASS and various control groups lacking components of the COMPASS. w/o, without. Scale bar, 100 μm. **b,** Flow cytometric analysis of COMPASS-processed MCF-7 cells incubated with Ac_4_ManNAz, 5-EU, or both, followed by sequential treatment with DBCO-AF647 and N_3_-AF488. **c,** RT-qPCR analysis of RNA samples extracted from *DTWD2* gene silencing by siRNA and the negative control (NC). **d,** CLSM images of MCF-7 cells after silencing of the *DTWD2* gene by siRNA or NC. Scale bar, 20 μm. **e,** Quantification of average fluorescence intensity in **d**. Data in **e** is representative of three independent experiments; n = 5 frames. **f,** Colocalization images (overlay channel) of glycoRNAs (COMPASS channel) with plasma membrane (DiD channel) or lipid rafts (CT-B channel). Scale bar, 10 μm. The line diagram depicts the intensity profiles of the fluorescent signals along the yellow arrows in the overlay images. The Pearson scatterplot on the right illustrates the degree of colocalization in the overlay image. Data are shown as mean ± s.d. Statistical significance was analyzed by unpaired t-test; NS, not significant;(*) P < 0.05, (**) P < 0.01, (***) P < 0.001, and (****) P < 0.0001.

We further optimized the MCR incubation time based on FRET signal intensity (**Supplementary Fig. 7**), along with the conditions for the subsequent click reactions, including the concentration and incubation time of DBCO–AF647 (**Supplementary Figs. 8 and 9**) and N_3_–AF488 (**Supplementary Figs. 10 and 11**). Under these optimized conditions, we confirmed that N_3_ groups on glycans were efficiently conjugated to DBCO–AF647, leaving undetectable residual N_3_ available to cross-react with EU-labeled RNA during the CuAAC reaction (**Supplementary Fig. 12**). Under optimized conditions, flow cytometric analysis of MCF-7 cells revealed a substantial increase in the FRET-positive population following COMPASS labeling, 11.5-fold higher than that observed in cells labeled for RNA alone, and 35.9-fold higher than in cells labeled for glycans alone (**Fig. 2b**).

The *DTWD2* gene was identified to regulate the assembly of glycans to RNAs, with its loss leading to the downregulation of glycoRNA levels^11^. To validate that the FRET signal arises from glycans and RNAs located in the same glycoRNA molecule, we established a gene silencing model using small interfering RNA (siRNA) to specifically knock down *DTWD2* expression. Efficient *DTWD2* knockdown in MCF-7 cells was confirmed by monitoring siRNA internalization (**Supplementary Fig. 13**) and gene expression levels (**Fig. 2c**), with no significant impact on cell viability (**Supplementary Fig. 14**). Notably, COMPASS imaging revealed a substantial reduction in FRET signal following *DTWD2* knockdown (**Figs. 2d and 2e**), while the individual AF488 and AF647 signals remained largely unchanged. Given that *DTWD2* knockdown specifically downregulates glycoRNA expression, the observed decrease in FRET signal strongly supports that the COMPASS-detected FRET arises from intact glycoRNA molecules.

The three-dimensional reconstructed image of a single MCF-7 cell captured with the assistance of COMPASS shows that the majority of FRET signals were localized at the cell membrane surrounding the nucleus (**Supplementary Fig. 15**). Further investigation into the subcellular localization of glycoRNAs revealed a concordance between COMPASS-derived FRET signals and those of cell membrane probe DiD and lipid raft probe CT-B (**Fig. 2f)**, with Pearson’s coefficients of 0.7601 and 0.7550, respectively. This relationship was further validated by z-stack images and orthographic projections (**Supplementary Figs. 16 and 17**). These results indicate that the COMPASS-derived FRET signals are predominantly situated on lipid rafts within the plasma membrane, consistent with the spatial distribution of glycoRNAs^8, 14^, thereby reinforcing the feasibility of COMPASS for glycoRNA imaging.

### Assessing COMPASS specificity and generality

The specificity of COMPASS for glycoRNA imaging was assessed by treating MCF-7 cells with ribonucleases (RNases), glycosidases, glycosylation inhibitors, and proteases. Treatment with RNase A and RNase T1 significantly reduced AF488 signals, while AF647 signals remained largely unaffected (**Figs. 3a and 3b**), consistent with selective degradation of cell surface RNAs and indicating that glycans on RNA contribute minimally to the overall glycan profile. In parallel, FRET signals were reduced by 67.74% and 69.37% following RNase A and RNase T1 treatment, respectively, confirming that RNA is required for COMPASS to generate FRET signals. Next, we treated cells with PNGase F and α2-3,6,8 neuraminidase (NA), which cleave N-glycans and terminal sialic acids, respectively^11, 32^. These treatments decreased FRET signals by 66.45% and 67.94%, and abolished glycan signals (**Figs. 3a and 3b**), indicating that N-glycans and sialic acids are also essential for effective COMPASS imaging of glycoRNAs. Then, we disrupted glycoRNA biosynthesis with glycosylation inhibitors before COMPASS labeling^33, 34^. The N-glycosylation inhibitor tunicamycin reduced FRET signals by 60.54%, while the O-glycosylation inhibitor benzyl-N-acetyl-α-d-galactosaminide (BG) had only a minor effect (**Figs. 3a and 3b)**, consistent with the predominantly N-glycan nature of glycoRNAs^8^. Furthermore, given the recent discovery that glycoRNAs can associate with glycosylated RNA-binding proteins (RBPs) at the cell surface^35^, we examined whether RBP labeling might interfere with FRET signals. Efficient protease digestion of RBPs, including trypsin and proteinase K (**Supplementary Fig. 18**), significantly reduced glycan signals while causing limited influence on FRET signals (**Figs. 3a and 3b**), indicating that COMPASS signals are derived from glycoRNA instead of RBP–RNA complexes. Additionally, a COMPASS-like immunofluorescence approach targeting cell surface RBPs failed to produce detectable FRET signals (**Supplementary Fig. 19**), suggesting that the distance between RBPs and RNA, although modestly increased by the use of antibodies, exceeds the effective FRET range and remains greater than that between the glycan and RNA within a single glycoRNA molecule. Together, these results demonstrate that COMPASS specifically reflects glycoRNA localization.

**Fig. 3.**
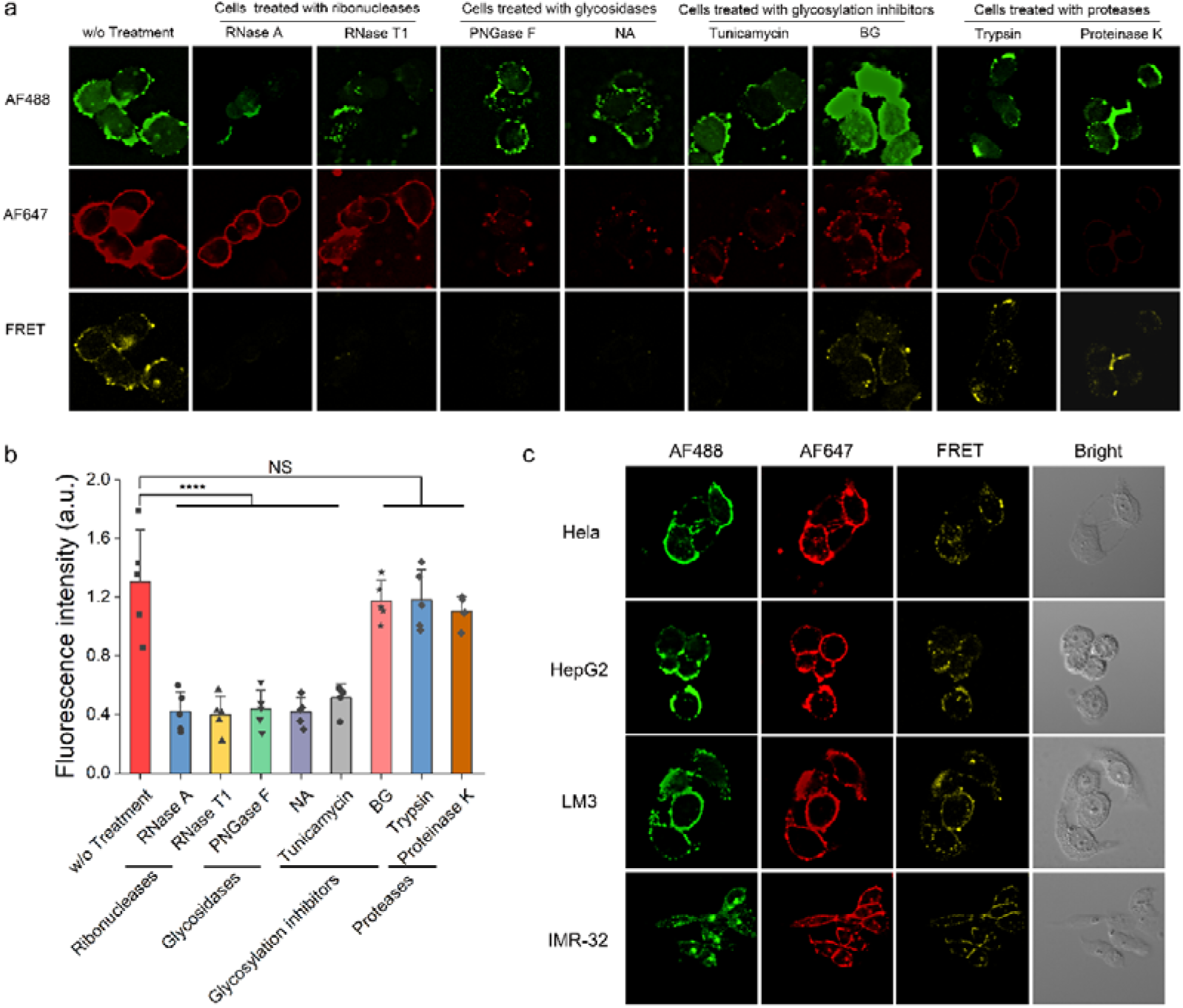
Evaluation of the specificity and generality of COMPASS in imaging glycoRNAs in cells. **a,** Specificity for COMPASS to image glycoRNAs on the MCF-7 cell surface. Imaging with COMPASS after treating MCF-7 cells with two RNases (RNase A and RNase T1), two glycosidases (PNGase F and NA), two glycosylation inhibitors (tunicamycin and BG), or two proteases (trypsin and proteinase K), respectively. Scale bar, 20 μm. **b,** Quantification of average fluorescence intensity in **a**. Data in **b** is representative of three independent experiments; n = 5 frames. Data are shown as mean ± s.d. Statistical significance was analyzed by unpaired two-tailed Student’s t-test; NS, not significant;(*) P < 0.05, (**) P < 0.01, (***) P < 0.001, and (****) P < 0.0001. **c,** Generality for COMPASS to image glycoRNAs on the cell surface of Hela cells, HepG2 cells, LM3 cells, and IMR-32 cells. Scale bar, 20 μm.

To evaluate the generality of COMPASS, we expanded its use to image various cell lines, including a human cervical carcinoma cell line (Hela), a human hepatocellular carcinoma cell line (HepG2), a human high metastatic potential hepatocellular carcinoma cell line (LM3), and a human neuroblastoma cell line (IMR-32). COMPASS-enabled CLSM imaging showed intensified signals corresponding to RNAs, glycans, and glycoRNAs on all the cell lines (**Fig. 3c**). Interestingly, the presence of glycoRNAs in the neuronal-like cell line (IMR-32) was observed for the first time, and further investigation revealed the appearance of glycoRNAs in a rat adrenal pheochromocytoma cell line (PC-12) (**Supplementary Fig. 20**), a well-established cell model in neurobiology research^36^. The imaging results from multiple cell lines underscore the broad applicability of COMPASS for visualizing cell surface glycoRNAs.

### Visualization of glycoRNAs in breast cancer development

Abnormal glycosylation is related to tumor development and progression^37–39^, particularly through altered sialic acid patterns that promote metastasis^40^. GlycoRNAs are characterized by containing sialic acid-terminated glycans, they are likely involved in cancer biology, though their roles remain poorly defined. To explore this, we utilized COMPASS to image glycoRNAs on non-tumorigenic (MCF-10A) and malignant (MCF-7) breast cancer cell lines. Comparable AF488 signals were observed between the two cell types, while AF647 signals were elevated in MCF-7 cells (**Fig. 4a and Supplementary Fig. 21**), consistent with the positive correlation between cell surface sialylation and malignancy. However, strikingly, the FRET signal in MCF-10A cells was threefold higher than that in MCF-7 cells (**Figs. 4a and 4b**). Although differences in MCR labeling efficiency between the two cell lines could potentially influence FRET signals, the higher AF647 (acceptor) and stable AF488 (donor) signals in MCF-7 cells would normally predict an equal or stronger FRET signal. The observed reduction instead suggests an inverse relationship between glycoRNA abundance and malignancy. This trend is consistent with recent findings showing downregulation of three sequence-specific glycoRNAs (U1, U35a, Y5) in malignant cells^8^. Our results suggest that this inverse relationship may extend more broadly across glycoRNAs, independent of RNA sequence.

**Fig. 4.**
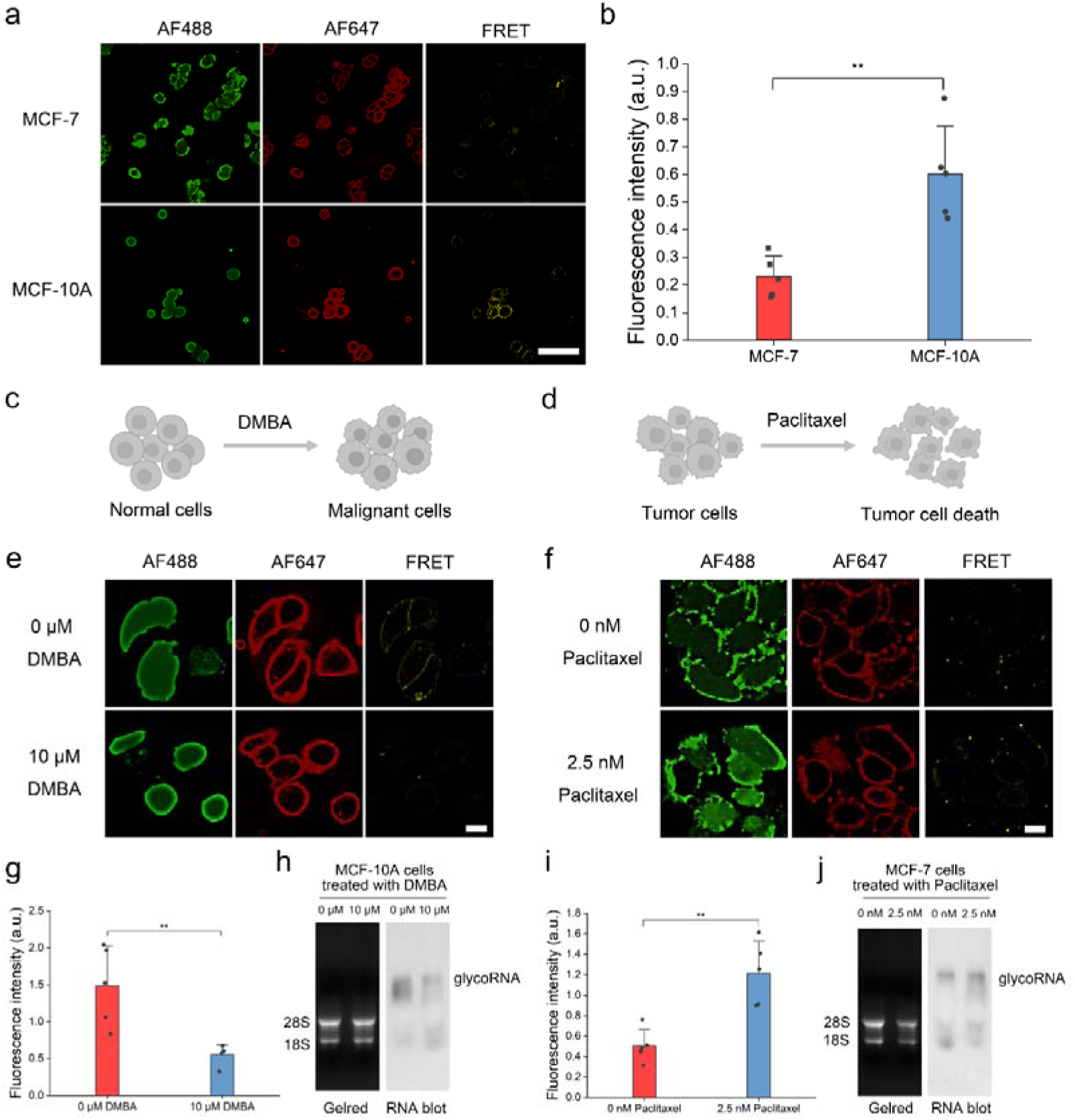
Imaging of glycoRNAs in cancer cell development. **a,** CLSM images of glycoRNAs by COMPASS in MCF-7 and MCF-10A cells. Scale bar, 100 μm. **b,** Quantification of the average fluorescence intensity of the FRET channel in **a**. Data in **b** is representative of three independent experiments; n = 5 frames. Data are shown as mean ± s.d. **c,** Schematic illustration of the malignancy transformation model, in which MCF-10A cells were exposed to DMBA to induce malignant transformation. **d,** Schematic illustration of chemotherapy model, in which MCF-7 cells were treated with PTX to induce cell death. **e,** Visualization of glycoRNAs abundance after treating MCF-10A cells with DMBA. Scale bar, 20 μm. **f,** Visualization of glycoRNAs after treating MCF-7 cells with PTX. Scale bar, 20 μm. **g,** Quantification of the average fluorescence intensity of the FRET channel in **e**. Data in **g** is representative of three independent experiments; n = 5 frames. Data are shown as mean ± s.d. **h,** The expression levels of total glycoRNA in MCF-10A cells treated with different concentrations of DMBA were assessed by MCR-free RNA blotting. **i,** Quantification of the average fluorescence intensity of the FRET channel in **f**. Data in **i** is representative of three independent experiments; n = 5 frames. Data are shown as mean ± s.d. **j,** The expression levels of total glycoRNA in MCF-7 cells treated with different concentrations of PTX were assessed by MCR-free RNA blotting. Statistical significance was analyzed by unpaired t-test; NS, not significant;(*) P < 0.05, (**) P < 0.01, (***) P < 0.001, and (****) P < 0.0001.

To investigate whether glycoRNA expression fluctuates during malignant transformation and in response to therapeutic intervention, we established two cellular models. A malignancy transformation model was generated by treating non-tumorigenic MCF-10A cells with 10 μM DMBA (**Fig. 4c and Supplementary Fig. 22**), a carcinogen known to induce epithelial cell transformation^41, 42^. Parallelly, a chemotherapy model was established by treating malignant MCF-7 cells with 2.5CnM paclitaxel (PTX) (**Fig. 4d and Supplementary Fig. 26**), an antineoplastic agent known to induce apoptosis^43^.

In the DMBA-induced transformation model, wound healing and RT-qPCR assays confirmed malignant characteristics in MCF-10A cells, including enhanced migratory capacity, upregulation of *EGFR*, and downregulation of the tumor suppressor *p53* (**Supplementary Figs. 23 and 24**). COMPASS imaging revealed that DMBA-treated cells exhibited similar AF488 and AF647 signals compared to controls (**Fig.**0**4e and Supplementary Fig.**0**25**), but a 62.89% reduction in the FRET signal (**Figs.**0**4e and 4g**). Considering the potential interference of DMBA with MCR labeling efficiency, an RNA blotting analysis independent of MCR incorporation was performed, which further confirmed this trend (**Fig. 4h**). These results suggest that downregulation of glycoRNAs is associated with early oncogenic transformation.

In the chemotherapy model, downregulation of *EGFR* and upregulation of *p53* were observed (**Supplementary Fig.**0**27**). Notably, the FRET signal increased by 58.57% compared to untreated controls (**Fig.**0**4f and 4i**), while AF488 signals remained comparable and AF647 signals decreased slightly (**Supplementary Fig.**0**28**). The increased expression of glycoRNA was further confirmed by MCR-independent RNA blotting (**Fig.**0**4j**), indicating that PTX-induced cell death correlates with elevated glycoRNA levels. Together, these findings demonstrate that cell surface glycoRNAs are dynamically regulated in response to malignancy transformation and chemotherapeutic stress, supporting their potential as responsive biomarkers in cancer biology.

### COMPASS-assisted glycoRNA imaging in organoids and tissues

The robust cellular applicability of COMPASS prompted us to extend its utility to more complex biological systems, including three-dimensional organoids and tissues. Building on our observation of glycoRNAs in neuronal cell lines (IMR-32 and PC-12), we applied COMPASS to cerebral organoids derived from human pluripotent stem cells (hPSCs) (**Fig. 5a and Supplementary Fig. 29**), which are widely used for modeling human brain development and neurological disorders^44, 45^.

**Fig. 5.**
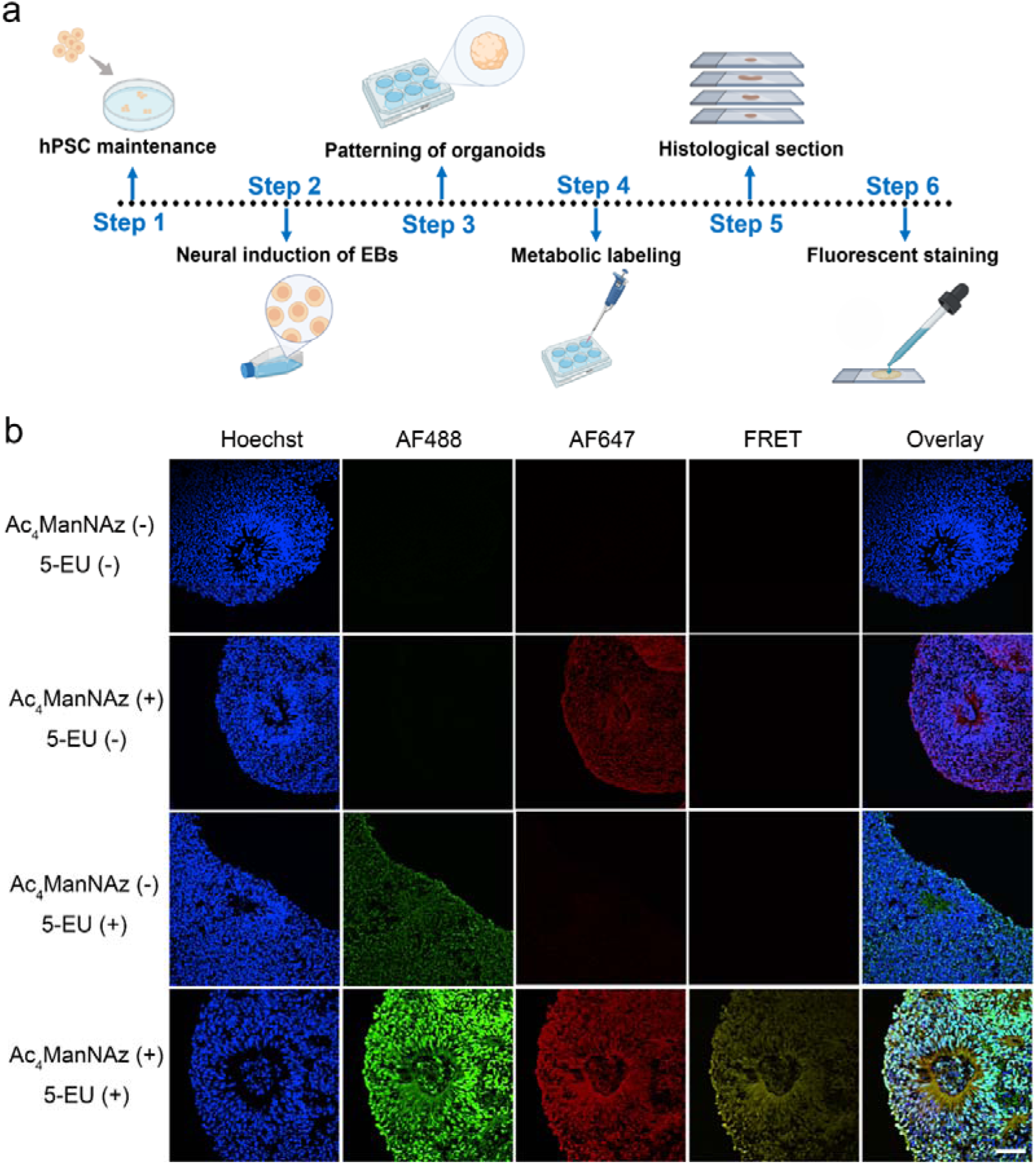
Visualization of glycoRNAs distribution in human cerebral organoid. **a,** Schematic illustration of the experimental procedure for culturing cerebral organoids and performing COMPASS-assisted imaging of glycoRNAs within the cerebral organoids. **b,** Visualization of glycoRNAs in human cerebral organoids using COMPASS. Human cerebral organoids were incubated with single or both MCRs (50 μM Ac_4_ManNAz, 250 μM 5-EU) in neural induction medium for 48 h. Subsequently, all organoid sections were stained for AF647 and AF488 by a two-step click chemistry reaction. Scale bar, 50 µm.

CLSM revealed well-organized radial arrangements of neural progenitors forming a neural tube-like architecture within the organoids (**Fig. 5b**). We established four experimental groups, all of which were co-stained with N_3_-AF488 and DBCO-AF647 but differed in their exposure to the 5-EU and Ac_4_ManNAz. AF488 fluorescence was observed exclusively in groups treated with 5-EU, while AF647 signals appeared only in groups exposed to Ac_4_ManNAz. Notably, only the group co-treated with both MCRs exhibited concurrent AF488, AF647, and FRET signals (**Fig. 5b**), confirming the specificity of COMPASS and minimal nonspecific dye adsorption in cerebral organoids. Of particular interest, the apical surface of the organoids, corresponding to the ventricular zone (VZ)-like region, exhibited elevated glycan and glycoRNA signals. The VZ-like zone serves as a critical neurodevelopmental niche, enriched in neural stem cells responsible for generating neurons and intermediate progenitor cells^46, 47^. The enrichment of glycan and glycoRNA signals in this region suggests that both general glycosylation and RNA glycosylation are heightened in neural stem cells with high differentiation potential, implicating the association between glycoRNA and neurogenesis in early brain development.

We next employed COMPASS to investigate glycoRNA distribution in animal tissues using a well-established 4T1 breast cancer metastasis mouse model^48^. GlycoRNA expression in the 4T1 cell line was first validated via COMPASS-assisted imaging (**Supplementary Fig. 30**). The incorporation of 5-EU and Ac_4_ManNAz into glycoRNAs in mouse tissues was further confirmed by enriching tissue-derived glycoRNAs using DBCO-functionalized magnetic beads, followed by RNA release, hydrolysis to nucleosides, and HPLC analysis (**Supplementary Fig. 31a**). Chromatographic profiles of glycoRNA-derived hydrolyzed nucleosides from major tissues revealed clear 5-EU signals by comparison with authentic standards (**Supplementary Figs. 31b and 31c**), demonstrating the in vivo robustness of co-metabolic labeling of these two MCRs to glycoRNAs.

Mice in the model group were then injected with 4T1 cells, while control animals received blank buffer. Both groups were administered injections of 5-EU and Ac_4_ManNAz. Three hours before sacrifice, DBCO-AF647 was injected, followed by tissue harvesting and a CuAAC reaction to conjugate N_3_-AF488 to RNA (**Fig. 6a**). Prominent nodules were observed on the lung surface of 4T1-injected mice (**Fig. 6b**), and H&E staining of major organs confirmed the presence of pulmonary metastases in the model group (**Fig. 6c and Supplementary Fig. 32**), validating the establishment of the metastatic model.

**Fig. 6.**
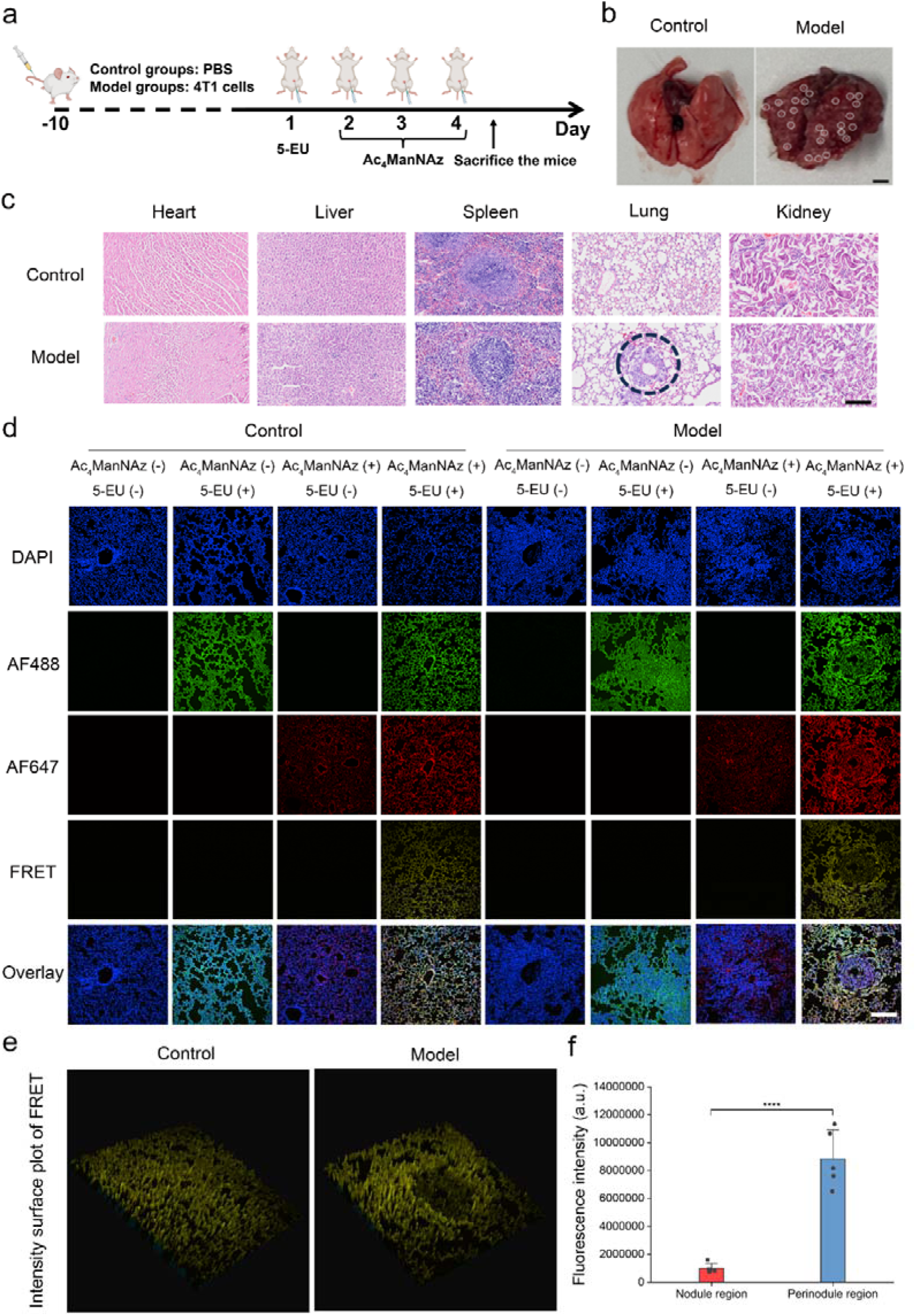
Visualization of glycoRNAs distribution in tissues of the mouse breast tumor metastasis model. **a,** Schematic illustration of the experimental procedure for imaging mouse tissues using COMPASS. Balb/c mice (n=3) were i.v. injected with 4T1 cells in the model groups, while PBS was injected in the control group. The 5-EU (20 mg/ml) was i.p. injected initially, followed by Ac_4_ManNAz (5 mg/kg) once daily for the following three days. **b,** Images of the lung harvested from mice. Metastatic nodules in the model group are indicated by black circles. Scale bar, 2 mm. **c,** Hematoxylin & eosin (H&E) staining of tissues from the major organs of control and model mice. Metastatic nodules in the model group are indicated by black circles. Scale bar, 100 µm. **d,** CLSM images of FRET signals in lung tissues from different treated mice captured with the assistance of COMPASS. The experimental mice were divided into the following two groups (n = 3 per group): a control group and the model group. To establish tumor metastasis models, 4T1 cells (1×10^7^ cells) suspended in PBS were injected intravenously through the tail vein in the model group. Meanwhile, the control group was injected with PBS. Mice were labeled with a single or two MCRs (100 μl of 20 mg/ml 5-EU, 0.16 mmole/kg DBCO-AF647), respectively, and subsequent slices were subjected to two-step click chemistry to achieve AF488 and AF647 labeling. Scale bar, 100 µm. **e,** Intensity surface plots of FRET signals in lung tissues from different treated mice captured with the assistance of COMPASS. **f,** Quantification of the average fluorescence intensity of nodule versus perinodule regions in the FRET channel on the right side of **e**. Data in **f** is representative of three independent experiments; n = 5 frames. Data are shown as mean ± s.d. Statistical significance was analyzed by unpaired t-test; NS, not significant;(*) P < 0.05, (**) P < 0.01, (***) P < 0.001, and (****) P < 0.0001.

To validate the application of COMPASS in tissue section imaging, we performed a panel of control conditions in both groups by modulating the administration of 5-EU and Ac_4_ManNAz while consistently applying both N_3_-AF488 and DBCO-AF647. AF488 signals were observed only in 5-EU-treated ones, AF647 signals only in Ac_4_ManNAz-treated ones, and FRET signals appeared exclusively in samples treated with both MCRs (**Fig. 6d and Supplementary Figs. 33–36**), demonstrating the feasibility and specificity of COMPASS for imaging glycoRNAs in tissue.

Notably, in lung sections from tumor-bearing mice, significantly lower FRET signals were detected within metastatic nodules compared to surrounding alveolar and perinodular regions (**Fig. 6d**). Surface plots of FRET intensities revealed pronounced signal depressions in nodule regions of the model group that were absent in controls (**Fig. 6e**), and quantitative analysis confirmed a substantial reduction in FRET signal that corresponding to glycoRNA abundance within the nodule region (**Fig. 6f**). This observation is consistent with the inverse correlation between glycoRNA levels and malignancy seen in vitro (**Fig. 4**), now extended to tissue-level validation. Furthermore, COMPASS enabled the profiling of glycoRNA distribution across tissue substructures (**Supplementary Figs. 33–36**). For instance, glycoRNA expression was significantly lower in renal glomeruli than in renal tubules (**Supplementary Fig. 36**), indicating heterogeneous glycoRNA distribution linked to cell-type-specific functions. Together, these findings underscore the capability of COMPASS for spatially resolved glycoRNA imaging in tissues and support its potential application in preclinical and translational research.

## Discussion

Here, we present COMPASS, a sequence-independent and in situ imaging strategy for glycoRNAs. Co-metabolic incorporation of two MCRs was validated at both cellular and in vivo levels by enriching glycoRNAs with DBCO-functionalized MBs, followed by RNA release, hydrolysis to nucleosides, and HPLC detection of 5-EU signals (**Figs. 1f–k and Supplementary Fig. 31**). HPLC analyses further confirmed minimal metabolic interference between the two reporters (**Supplementary Figs. 4 and 5**). The specificity of COMPASS for glycoRNAs was established through selective perturbations using RNases, glycosidases, proteases, and glycosylation inhibitors (**Figs. 3a and 3b**). Importantly, COMPASS enabled direct visualization of glycoRNAs across diverse biological systems, including multiple cell lines (**Figs. 3c and 4; Supplementary Figs. 20 and 30**), human cerebral organoids (**Fig. 5**), and mouse tissues (**Fig. 6 and Supplementary Figs. 33–36**).

Given that glycans are broadly distributed on the cell surface, FRET arising from spatial proximity between RNA and separated, neighboring glycans could theoretically contribute to background signal^7^. However, the effective FRET distance (∼10 nm) imposes a physical constraint, such that this background is dependent on the glycan density within a 10 nm radius of labeled RNAs. In a previous study utilizing a 14.3 nm DNA connector to bridge glycan and RNA probes, background signal arising from separate glycans was found to be negligible^8^, suggesting that the spatial co-localization of separated glycans within 10 nm of RNA is rare. Importantly, knockdown of *DTWD2*, a gene recently implicated in glycan-RNA assembly, resulted in a marked reduction in FRET signal, with minimal changes in overall RNA or glycan labeling (**Figs. 2c–2e; Supplementary Figs. 13 and 14**). These findings provide direct evidence that COMPASS exhibits high specificity for intact glycoRNA molecules, with minimal contribution from non-covalently associated glycans.

The inverse relationship between glycoRNA expression and cell malignancy has been previously reported^8^, and this trend was recapitulated using COMPASS in our study (**Figs. 4a and 4b**). Extending this observation, we observed a pronounced reduction in glycoRNA expression during DMBA-induced transformation of MCF-10A cells, as evidenced by both MCR-incorporated COMPASS imaging and MCR-free RNA blotting (**Figs. 4e, 4g, and 4h**). These findings suggest that glycoRNA downregulation may represent an early event in carcinogenesis. Moreover, in a mouse 4T1 breast cancer metastasis model, glycoRNA expression was significantly reduced within tumor nodules compared to perinodule regions, representing the first demonstration of glycoRNA downregulation in tumor progression at the tissue level.

While the widespread distribution of glycoRNAs has been established, their imaging in neuronal cells and human cerebral organoids has not been documented. We employed COMPASS to visualize glycoRNAs in human cerebral organoids, revealing distinct glycoRNA signals within the rosette-shaped structures of these organoids. Notably, although the MCRs were incubated with the organoids and penetrated from the outer to the inner layers, we observed an enhanced abundance of glycoRNAs in the inner layer, particularly on the apical surface of the VZ-like region. Since the VZ-like region is crucial for human brain development and contains cells with stem cell activity^44, 49^, the elevated glycoRNA abundance in this area suggests an association between glycoRNA and neurogenesis during early brain development.

Compared to existing methods for the cell imaging of glycoRNAs, such as ARPLA and HieCo-2, COMPASS offers several distinct differences and advantages (**Supplementary Table 1**). By labeling the RNA moiety through metabolic incorporation, COMPASS enables sequence-independent visualization of glycoRNAs, overcoming the requirement for prior sequence knowledge inherent to probe-based strategies. The method relies exclusively on commercially available MCRs and robust bioorthogonal click chemistry, eliminating the need for custom DNA probes or labile enzymatic reactions and substantially simplifying the workflow, with the time from completion of cell culture to imaging reduced to approximately 1 h compared with at least 4.5 h in previous methods. COMPASS further employs FRET as the signal readout, allowing channel-selective visualization of glycans, RNAs, and glycoRNAs while benefiting from reduced background owing to the large excitation–emission separation intrinsic to FRET. Direct labeling with short chemical linkers also improves spatial fidelity by avoiding the signal displacement associated with long DNA amplification products ^16, 17^. Importantly, COMPASS shows broad compatibility across biological contexts, enabling glycoRNA imaging in cultured cells, organoids, and mammalian tissues, and providing the first spatial mapping of glycoRNA distribution in vivo. In light of recent reports identifying glycoRNAs in extracellular vesicles^15, 50^, this strategy may be readily extended to sequence-independent imaging of glycoRNAs in extracellular compartments, further broadening its applicability.

Despite its competitive performance in glycoRNA imaging, COMPASS has certain limitations. First, although its applicability to organoids and tissues was demonstrated, protocol optimization and standardization remain necessary to enhance reproducibility and cross-study comparability. Second, because COMPASS relies on direct FRET without signal amplification, its sensitivity is limited, particularly in detecting glycoRNAs, which are often present at low abundance. Integration with amplification, high-efficiency FRET systems could address this challenge^51^. Third, although sequence-independent, COMPASS currently detects glycoRNAs that (i) contain U and (ii) are sialylated. Rare glycoRNAs lacking U or possessing U too distant from the glycosylation site, as well as non-sialoglycoRNAs, may require alternative metabolic labeling or chemoenzymatic strategies^52, 53^. Fourth, metabolic labeling targets only newly synthesized glycoRNAs, leaving pre-existing glycoRNAs undetectable; engineering of proximity labeling techniques could provide a route to visualize the intrinsic pool efficiently^54, 55^. Lastly, the CuAAC-based dye conjugation step restricts COMPASS to fixed samples and precludes real-time in vivo applications. Moreover, the reliance on MCRs without clinical approval poses a significant barrier to clinical translation. Future advances will require the development of biocompatible labeling agents with regulatory approval potential and the implementation of label-free sensing platforms capable of generating direct on/off signals upon glycoRNA recognition, thereby enabling translational applications of glycoRNA imaging.

## Acknowledgments

This research was supported by the Jiangsu Basic Research Center for Synthetic Biology (BK20233003), the National Natural Science Foundation of China (22304083 and 22274079), and the Natural Science Foundation of the Jiangsu Higher Education Institutions of China (25KJB330004). We thank Prof. Yan Liu at the School of Pharmacy at Nanjing Medical University for providing the conditions to conduct experiments regarding human cerebral organoids, Prof. Ran Xie at the School of Chemistry and Chemical Engineering at Nanjing University for providing Neu5Az, and Dr. Furong Zhao at Southeast University for providing the 4T1 cell line. Confocal imaging was performed at the Analytical and Testing Center of Nanjing Normal University. We thank Prof. Lei Lin at Nanjing Normal University for providing advice on confocal imaging and result interpretation. We thank Dr. Jie Zheng, Dr. Kaiheng Fang, Dr. Xiaolong Zheng, and Mr. Linfang Wu for their valuable academic suggestions, manuscript proofreading, and technical support with RNA blotting.

## Author contributions

B.L., X.Z., and H.H. conceived and designed the study. B.L. and S.L. performed the experiments and analyzed the data. S.W. assisted in validating the COMPASS and culturing cells. X.L. performed RNA blotting experiments. Y.C. designed the cancer cell-related investigations and analyzed the data. X.Z. performed the MD simulations. R.J. assisted in click chemistry reactions. W.Z. and Y.F. performed human cerebral organoid experiments. The manuscript was written by B.L., S.L., and X.Z. H.H. supervised the project and provided the funding support.

## Methods

### Materials

Ac_4_ManNAz was purchased from Jinan Samuel Pharmaceutical Co., Ltd. DBCO-AF647 was obtained from Confluore. BeyoClick™ EU RNA Synthesis Kit with Alexa Fluor 488, RNase T1, PNGase F, Cell Plasma Membrane Staining Kit with DiD, and 0.45-µm nitrocellulose (NC) membrane were supplied by Beyotime Biotechnology. Antibodies, including anti-hnRNP U antibody (Cat No. 14599-1-AP) and CoraLite647-conjugated F(ab’)2 Fragment Goat Anti-Rabbit IgG (H+L) (Cat No. SA00014-9) were obtained from Proteintech. α2-3,6,8,9 Neuraminidase A (NA) was purchased from New England Biolabs (NEB). RNase A, RNATransMate, and siRNA were provided by Sangon Biotech. Tunicamycin (TM) and Benzyl-2-acetamido-2-deoxy-α-D-galactopyranoside (BG) were provided by Macklin. AF594-labeled cholera toxin subunit B (CT-B) was acquired from Thermo Fisher Scientific. TransZol Up was purchased from TransGen Biotech. Paclitaxel and 7,12-dimethylbenzanthracene (DMBA) were purchased from MedChemExpress (MCE).

### Cell Culture

MCF-7 cell line, MCF-10A cell line, Hela cell line, HepG2 cell line, PC-12 cell line, and IMR-32 cell line were derived from ATCC. The 4T1 cell line was a gift from Dr. Furong Zhao at Southeast University. The LM3 cell line was obtained from the National Collection of Authenticated Cell Cultures. All cell lines were cultured at 37C°C in a humidified incubator with 5% CO_2_.

Breast cancer cell line MCF-7 (HTB-22, ATCC) and non-tumorigenic breast cell line MCF-10A (CRL-10317, ATCC) were cultured in RPMI 1640 medium supplemented with 10% fetal bovine serum (FBS, Tianhang), 100 μg/ml penicillin, and 100 μg/ml streptomycin.

Hela cell line (CRM-CCL-2, ATCC), HepG2 cell line (HB-8065, ATCC), 4T1 cell line (CRL-2539, ATCC), LM3 cell line (SCSP-5093), and PC-12 cell line (CRL-1721, ATCC) were treated in DMEM supplemented with 10% FBS, 100 μg/ml penicillin, 100 μg/ml streptomycin, and 4.5g L^-1^ D-glucose. IMR-32 cell line (ATCC, CCL-127) was cultured in the MEM medium supplemented with 10% FBS and 100 μg/ml penicillin and streptomycin.

### Blotting analysis of glycoRNA with metabolic labeling

GlycoRNAs from MCF-7 cells were metabolically labeled with Ac_4_ManNAz, and then the presence of glycoRNAs was confirmed by RNA blotting, as previously described^4^. Subsequently, total RNA was extracted and purified. DBCO-biotin was labeled on the purified RNA using biorthogonal click chemistry and then analyzed by denaturing gel electrophoresis and RNA blotting. As shown in **Fig. S2**, glycoRNAs were detected in total RNA extracted from metabolically labeled MCF-7 cells compared to MCF-7 cells without metabolic labeling treatment, demonstrating the presence of glycoRNAs in MCF-7 cells.

#### Metabolic labeling

MCF-7 cells were seeded in T75 cell culture flasks and cultured for 24 h, followed by treatment with 100 µM Ac_4_ManNAz in the cell culture medium for 72Ch.

#### RNA extraction and purification

Firstly, the cells were treated with 6 ml of TRIzol reagent and incubated at 37 °C. After denaturation, the phases were separated by adding 0.4 ml of chloroform, vortexing, incubating for 5 min, and centrifuging at 12,000 g for 15 min at 4 °C. The aqueous phase was transferred to a new centrifuge tube, mixed upside down with isopropanol, incubated for 10 min at room temperature, and then centrifuged at 10,000g for 10 min at 4°C. After removing the supernatant, 1 ml of 75% ethanol was added, vortexed vigorously, and centrifuged at 7500 g for 5 min to obtain RNA precipitate.

#### Copper-free click conjugation to RNA

To perform the SPAAC, Ac_4_ManNAz-labeled RNA (10µg) was mixed with RNA denature buffer (95% formamide, 18 mM EDTA, and 0.025% SDS) and 500 μM DBCO-biotin. The reaction was performed at 55 °C for 10 min. Biotin-labeled RNA was then purified with a Zymo RNA Clean and Concentrator 5 kit. The concentration was measured with a NanoDrop (Thermo Fisher Scientific) and stored at -80 C for future use.

#### RNA gel electrophoresis, blotting, and imaging

Blotting analysis of biotin-labeled RNA was performed conceptually similar to a previously reported method^4^. RNA (10 μg) was added to the 10 μl 2×loading buffer (Beyotime Biotechnology), incubated at 55°C for 10 min, and then cooled on ice. Afterward, samples were loaded into a 1% agarose-formaldehyde denaturing gel (1.5×MOPS buffer, 0.75% formaldehyde, 1% agarose, 10ul 4sGelRed staining) and electrophoresed at 150 V for 45 min. Meanwhile, the total RNA in the gel was observed with a UV gel imager. RNA transfer to a 0.45-µm NC membrane occurred in a transfer buffer (3M NaCl, pH=1) for 16 h at room temperature. Afterward, the RNA sample was cross-linked to the NC membrane with UV for 2 min. The NC membrane was then treated with a protein-free blocking solution (Sangon Biotech) and incubated for 45 min at 25 °C before staining with IRDye 800CW streptavidin (Kamoer) for 30 min. It is important to note that the IRDye 800CW streptavidin was diluted 10,000-fold in the protein-free blocking solution. To flush out the excess IRDye 800CW streptavidin, the NC membrane was washed three times with 1×PBS containing 0.1% Tween-20 and then once with 1× PBS. Images were obtained by scanning the NC membrane with a Chemi Dog 5200T automatic luminescence imaging system (Tanon), and the software was set to automatically detect the signal intensity of 790 nm channels.

### MCR-free blotting analysis of glycoRNA

MCR-free blotting analysis of glycoRNA was conducted according to a previous report with a few modifications^11^. MCF-10A cells were seeded in T75 cell culture flasks and cultured for 24 h, followed by treatment with blank buffer or 10 µM DMBA in the cell culture medium for 24 h. The cells were washed with PBS, and total RNA was isolated by using TRIzol reagent. A 3 μg RNA sample was mixed with 2× RNA loading buffer. Add 28 μL of Blocking Buffer to the lyophilized RNA, vortex to mix thoroughly, and incubate at 35°C for 45 minutes for blocking. After cooling to room temperature for 2-3 minutes, 1 μL of 30 mM aldehyde reaction probe was added, followed by 2 μL of fresh 7.5 mM NaIOC. The oxidation reaction was carried out in the dark at room temperature for a full 10 min. The reaction was then quenched with 3 μL of freshly prepared 22 mM sodium sulfite and incubated at 25 °C for 5 min. Samples were returned to 35 °C for 90 min for the linkage reaction. The mixture was purified using a Zymo-I column according to the standard Zymo protocol, first diluting with 19 μL of water to a total volume of 50 μL. To display the RNA on an agarose gel, the RNA was eluted from the column with two 6.2 μL water washes, resulting in a final eluate of approximately 12 μL. Denaturation involved incubating the RNA at 55 °C for 10 minutes, then quickly cooling it on ice for 3 min. The samples were then applied to a 0.7% agarose-formaldehyde denaturing gel containing 4S Gelred nucleic acid dye (10,000× aqueous solution, Sangon Biotech). Electrophoresis was performed at 150 V for 30 minutes. A JY04S-3E (Beijing Junyi Electrophoresis Co., Ltd) was used to visualize the total RNA in the gel. Then, RNA was transferred to a 0.45-µm NC membrane in a transfer buffer (3M NaCl, pH=1) for 16 h at room temperature. After transfer, UV-C light at 0.18 J/cm² was applied to cross-link the RNA to the NC membrane. The NC membrane was then blocked for 30 minutes at 25 °C with QuickBlock™ Blocking Buffer (Beyotime). Following blocking, the membrane was treated with HRP-Streptavidin (1:3000) at 25 °C for 30 minutes. A brief rinse with 0.1% Tween-20 in 1x PBS removed excess HRP-Streptavidin. BeyoECL Plus working solution (Beyotime) was evenly applied to the membrane and left for 1-2 minutes. The membrane was then removed, and the BeyoECL Plus working solution was discarded. Finally, chemiluminescence imaging was conducted using a Chemiluminescence gel imager (Tanon 5200).

For MCF-7 cells, a similar procedure was used, except that blank buffer or 2.5 nM PTX was added to the cell culture medium and treated for 48 hours.

### Incorporation rate of Ac_4_ManNAz-labeled cell surface sialic acids

Experiments were conducted according to a previous report with a few modifications^27^.

#### Cell fractionation

MCF-7 cells were seeded in T75 cell culture flasks and cultured for 24 h, followed by treatment with 50 µM Ac_4_ManNAz in the cell culture medium for 48 h. The cell was resuspended in lysis buffer (10 mM Tris-HCl (pH=7.9), 150 mM NaCl, 5 mM EDTA, 1 mM PMSF, 0.462 μM aprotinin, 3 μM leupeptin, and 4.36 μM pepstatin A). Cell lysis was performed by ultrasonic fragmentation on ice at 20% intensity, 3 times for 30 s each, with a 1-minute interval between each time on ice. The lysate was then centrifuged at 21130 g for 10 min at 4 °C. The supernatant was collected and subjected to a second centrifugation under the same conditions at 21130 g for 60 min at 4 °C. The resulting precipitate was resuspended in 30 μL of fresh lysis buffer and centrifuged again (21130 g, 60 min, 4 °C). The supernatant from the above two steps was mixed and freeze-dried.

#### Reduction step

The freeze-dried product was centrally accompanied with 0.2 M NaBH_4_ solution (pH=8.0, 65 μL) overnight to remove free sialic acid. Excess NaBH_4_ was subsequently removed with TFA and then concentrated by vacuum drying.

#### DMB labeling of sialic acids

Treated cells with cell fractionation and reduction step were incubated with 300 μL of 3 M acetic acid at 80 °C with 300 rpm for 2 h. After incubation, the samples were immediately cooled on ice for 10 min to halt the reaction. Subsequently, DMB labeling was performed with the Sialic Acid Fluorescence Labeling Kit (Takara) and analyzed by HPLC with fluorescence detection.

#### HPLC analysis of sialic acid

Sialic acid derivatives were analyzed on a Shimadzu LC40 HPLC instrument equipped with a fluorescence detector using an Agilent 5 Prep-C18 Scalar 250 × 4.6 mm column. Sialic acid derivatives were separated under the following conditions: 45 min 100% acetonitrile/methanol/water (9/7/84) solution at a flow rate of 0.9 ml/min. The resulting fluorescent sialic acid derivatives were analyzed by using a fluorescence detector (λ_ex_=373 nm, λ_em_=448 nm). Incorporation rate (IR) of Ac_4_ManNAz-labeled cell surface sialic acids based on integrated area: IR = A_Neu5Az_ × (A^Neu5Ac^+ A_Neu5Az_) ^-1^ × 100%.

### Incorporation rate of 5-EU-labeled cell surface nucleosides

Experiments were conducted according to a previous report with a few modifications^22^.

#### RNA Isolation and Hydrolysis

MCF-7 cells were seeded in T75 cell culture flasks and cultured for 24 h, followed by treatment with 250 µM 5-EU in the cell culture medium for 24 h. The cells were washed with PBS, and total RNA was isolated by using TRIzol reagent. Purified RNA samples (5-50 μg) were hydrolyzed to nucleoside monophosphates by incubation in 50 mM potassium hydroxide at 100 °C for 1 h. The reaction mixture was then neutralized and adjusted to a final concentration of 50 mM Tris (pH=8.0), 40 mM KCl, 60 mM NaCl, and 10 mM MgClC. To dephosphorylate nucleoside monophosphates, 30 units of Quick CIP (New England Biolabs) were added, followed by incubation at 37°C for 2 h. The mixture is then freeze-dried and de-enzymatized with 80% ethanol (vortexed for 10 min at 37 ° C). After centrifugation to remove insoluble debris, the supernatant was collected, dried again, and resuspended in water.

#### HPLC analysis of nucleosides

Nucleosides were analyzed on a Shimadzu LC40 HPLC instrument equipped with a diode array detector using an Agilent 5 Prep-C18 Scalar 250 × 4.6 mm column. Nucleosides were separated under the following conditions: 30 min 5% acetonitrile in water at a flow rate of 0.4 ml/min. Pure standards for adenosine, cytidine, guanosine, and uridine were from Sangon. 5-EU standard was obtained from Beyotime. Nucleoside separation was monitored by absorbance at 260 nm. Incorporation rate (IR) of 5-EU was calculated based on integrated area: IR = A_5-EU_ × (A_U_+ A_5-EU_) ^-1^ × 100%.

### Confirming the co-incorporation two MCRs with glycoRNA enrichment and HPLC analysis

#### GlycoRNA enrichment and Hydrolysis

The culture of the MCF-7 cell line was performed as described above. Cells were divided into four groups: an unlabeled control maintained in standard culture medium; a 5-EU–labeled group cultured in medium supplemented with 5-EU at 250 μM; an Ac_4_ManNAz-labeled group cultured in medium supplemented with Ac_4_ManNAz at 50 μM; and a co-labeling group cultured in medium containing both 5-EU (250 μM) and Ac_4_ManNAz (50 μM). All treatments were carried out for 48 h.

Mouse handling and experimental procedures were performed as described in the section “Animal models and COMPASS-assisted glycoRNA imaging in mouse tissues.” Mice were randomly assigned to four groups (n = 3 per group). In the co-labeling group, mice received an intraperitoneal injection of 5-EU (100 μl, 20 mg/ml) on Day 1, followed by intraperitoneal injections of Ac_4_ManNAz (5 mg/kg) once daily for three consecutive days starting on Day 2. In the unlabeled control group, all injections were replaced with PBS. In the single-label groups, mice received either 5-EU or Ac_4_ManNAz alone at the same doses and on the same schedule as in the co-labeling group, with PBS administered in place of the omitted compound. All other housing conditions and experimental parameters were identical across groups.

After collecting cells and major organs (heart, liver, kidney, spleen, and lung), RNAzol® RT RNA Isolation Reagent (iGene Biotechnology) was used for small RNA extraction. After pipetting to ensure homogeneity, samples are incubated at room temperature for 15 minutes to further denature non-covalent interactions. The aqueous phase was carefully collected, mixed with 0.4 volumes of 75 % ethanol, and centrifuged at 10,000Crpm for 8 min at 4CC. The supernatant was transferred to a new tube, combined with 0.8 volumes of isopropanol, and incubated at -20CC for 30Cmin. After incubation, centrifuge at 4 °C at 10,000 rpm for 10 min. Wash the small RNA pellet twice with 400 μL of 75% isopropanol. After drying the small RNA pellet at room temperature, dissolve it in DEPC-treated water. Subsequently, the small RNA was treated with DNase I and proteinase K to remove DNA and protein contaminants.

We incubated 10 mg of DBCO-modified magnetic beads with small RNA at 37°C for 2 h to capture N_3_-labeled glycoRNA, followed by washing with buffer and cleavaging of the link between RNA and glycan using PNGase F. The supernatant was collected after a 4-hour reaction. RNA samples were hydrolyzed to nucleoside monophosphates by incubation in 50 mM potassium hydroxide at 100 °C for 1 h. The reaction mixture was then neutralized and adjusted to a final concentration of 50 mM Tris (pH=8.0), 40 mM KCl, 60 mM NaCl, and 10 mM MgCl_2_. To dephosphorylate nucleoside monophosphates, 30 U of Quick CIP (New England Biolabs) were added, followed by incubation at 37°C for 2 h. The mixture is then freeze-dried and de-enzymatized with 80% ethanol (vortexed for 10 min at 37 °C). After centrifugation to remove insoluble debris, the supernatant was collected, dried again, and resuspended in water.

#### HPLC analysis of nucleosides

Nucleosides were analyzed using a Shimadzu LC40 HPLC system equipped with a diode array detector. An Agilent Prep C18 column (250 × 4.6 mm) was used for samples derived from cultured cells, whereas a YMC Hydrosphere C18 column (250 × 4.6 mm) was used for animal tissue samples. Separations were performed with an isocratic mobile phase of 5% acetonitrile in water over 30 min at a flow rate of 0.4 ml/min. Authentic nucleoside standards of adenosine (A), cytidine (C), guanosine (G), and uridine (U) were obtained from Sangon, and the 5-EU standard was purchased from Beyotime. Nucleosides were detected by monitoring absorbance at 260 nm. IR of 5-EU into glycoRNA was calculated based on integrated area: IR = A_5-EU_ × (AU+ A_5-EU_) ^-1^ × 100%.

### In situ imaging of glycoRNAs with COMPASS

Taking MCF-7 as an example, cells were seeded at a density of 1.5×10^5^ cells per well on glass-bottom 35-mm imaging dishes (Nest) overnight. Before starting each step, cell samples were washed three times with PBS. Cells were incubated with Ac_4_ManNAz (50 μM) and 5-EU (250 μM) in culture medium for 48 h. After metabolic labeling, cells were incubated with 50 μΜ DBCO-AF647 for 10 min on ice. From this step onward, the well was kept in the dark. Then, the cells were treated with BeyoClick™ EU RNA Synthesis Kit with Alexa Fluor 488 (Beyotime Biotechnology) at 37 °C for 30 min. Finally, CLSM imaging of cells was acquired on an AX (Nikon). To accomplish the imaging, a ×40 water immersion objective was applied, and the RNA fluorescence signal (AF488 channel) was excited with a 488-nm laser and received at an emission of 500-550 nm. Glycan fluorescence signal (AF647 channel) was excited with a 640-nm laser and received at an emission of 662-737 nm. The GlycoRNA fluorescence signal (FRET channel) was excited with a 488-nm laser and received an emission of 662-737 nm.

The metabolic labeling durations and concentrations differ for each cell line, as detailed below. For HeLa cells, cells were incubated with Ac_4_ManNAz (50 μM) and 5-EU (250 μM) in the culture medium for 24 h. For HepG2 and LM3 cells, cells were incubated with Ac_4_ManNAz (50 μM) and 5-EU (100 μM) in culture medium for 48 h. For IMR-32, PC-12, and 4T1 cells, cells were incubated with Ac_4_ManNAz (50 μM) and 5-EU (250 μM) in culture medium for 48 h. For MCF-10A, cells were incubated with Ac_4_ManNAz (50 μM) and 5-EU (100 μM) in the culture medium for 24 h.

### Dynamic simulation of the glycoRNA nanostructure

To provide a theoretical basis for the application of COMPASS in imaging glycoRNA, we constructed a model of Y5 glycoRNA modified with AF647 using AlphaFold3^56^, replacing the U base in the RNA sequence with 5-EU. The complete 3D glycoRNA nanostructure is shown in **Fig. 1b**.

To further investigate the conformation and properties of the constructed model, as well as the distribution of U base around the AF647, we performed 100 ns kinetic simulations. MD simulations were performed using the Gromacs 2021.7 program^57^, applying the Amber99SB all-atom force field^58^, at constant temperature and pressure as well as periodic boundary conditions. The simulated system was solvated using the TIP3P water model^59^ in a cubic box where the minimum distance to the edge of the box was 2.0 nm, and counterions (Na^+^ and Cl^-^) were subsequently added to neutralize the charge of the system. Energy minimization was performed using the steepest descent with the maximum force set to 1000.0 kJ mol^-1^ nm^-1^ to eliminate too close contact between atoms. All bonds containing hydrogen atoms were constrained using the default Linear Constraint Solver (LINCS)^60^ algorithm. The V-rescale^61^ temperature coupling method was used to control the simulated temperature to 298.15 K, and the Parrinello-Rahman method^62^ was used to control the pressure to 1 bar. The electrostatic interactions were calculated using the (Particle-mesh Ewald) PME method^63^. Van der Waals interactions were calculated using a cutoff value of 14 Å. Then, NVT and NPT equilibrium simulations were performed at 298.15 K for 100 ps, respectively. Finally, the system was subjected to a 100 ns MD simulation with a time step of 2 fs, and conformations were saved every 50 ps. The visualization of the simulation results was done using the Gromacs embedded program and VMD.

### COMPASS-assisted flow cytometric analysis of glycoRNAs

MCF-7 cells were seeded on a six-well plate and cultured overnight. Before starting each step, cell samples were washed three times with PBS. Cells were incubated with Ac_4_ManNAz (50 μM) and 5-EU (250 μM) in culture medium for 48 h. After metabolic labeling, cells were incubated with 50 μΜ DBCO-AF647 for 10 min on ice. From this step onward, the well was kept in the dark. Then, the cells were treated with BeyoClick™ EU RNA Synthesis Kit with Alexa Fluor 488 at 37 °C for 30 min. Finally, the cells were dispersed in PBS for flow cytometric analysis (Beckman Coulter). Flow cytometric data were analyzed with FlowJo software. By contrast, for MCF-7 cells incubated with only Ac_4_ManNAz (50 μM) or 5-EU (250 μM) in the culture medium, no glycoRNA signal was observed (**Fig. 2b**).

### Validation of the specificity of COMPASS

MCF-7 cells were seeded at a density of 1.5×10^5^ cells per well on glass-bottom 35-mm imaging dishes overnight. After metabolic labeling with 50 μM Ac_4_ManNAz and 250 μM 5-EU in culture medium for 48 h, cells were incubated with 0.02 µg/µl RNase A or 1 U/µl RNase T1 at 37 °C for 30 min to digest the RNA moiety of glycoRNAs, respectively. Cells were subsequently treated with COMPASS.

Next, we removed glycan molecules using glycosidase and glycosylation inhibitors to validate the response of COMPASS to the glycan moiety in glycoRNAs. After metabolic labeling, MCF-7 cells were incubated with 0.1 U/µl PNGase F or 0.1 U/µl NA at 37 °C for 30 min and then analyzed by COMPASS.

In the glycosylation inhibitors treatment experiment, MCF-7 cells were incubated with 2 μΜ TM or 500 μΜ BG during metabolic labeling with 50 μM Ac_4_ManNAz and 250 μM 5-EU in culture medium. After 48 h of incubation, we stained and imaged using the method of COMPASS.

In trypsin and proteinase K treatment experiments, MCF-7 cells were incubated with 50 μM Ac_4_ManNAz and 250 μM 5-EU in culture medium. After 48 hours of culture, MCF-7 cells were incubated with trypsin or 0.02 mg/mL proteinase K at 37°C for 30 seconds. Floating cells were then gently washed away with PBS, and adherent cells were analyzed using the COMPASS system. ImageJ (Fiji) is used to calculate the average fluorescence intensity of the cells in each frame.

For RBP imaging, MCF-7 cells were first incubated with 2.5 mg/mL anti-hnRNP U antibody in an ice bath for 45 minutes. After washing, cells were labeled for 30 minutes with CoraLite 647-labeled F(ab’)2 fragment goat anti-rabbit IgG (H+L) secondary antibody. For the enzyme-treated control group, MCF-7 cells were incubated with trypsin or 0.02 mg/mL proteinase K at 37 °C for 30 seconds before the antibody and EU labeling procedures described above. Finally, CLSM imaging of cells was acquired on an AX (Nikon). To accomplish the imaging, a ×40 water immersion objective was applied, and the RBP fluorescence signal (CL647 channel) was excited with a 640-nm laser and received at an emission of 662-737 nm.

For a COMPASS-like procedure using immunofluorescence to label cell surface RBPs, MCF-7 cells were cultured for 48 hours in medium supplemented with 5-EU (250 μM). Cells were then incubated for 45 minutes with 2.5 mg/mL anti-hnRNP U antibody on ice, followed by incubation for 30 minutes with CoraLite647-labeled F(ab’)2 fragment goat anti-rabbit IgG (H+L) antibody. Subsequently, cells were treated with the BeyoClick™ EU RNA Synthesis Kit (containing Alexa Fluor 488) at 37°C for 30 minutes. Finally, CLSM imaging of cells was acquired on an AX (Nikon). To accomplish the imaging, a ×40 water immersion objective was applied, and the RNA fluorescence signal (AF488 channel) was excited with a 488-nm laser and received at an emission of 500-550 nm. RBP fluorescence signal (CL647 channel) was excited with a 640-nm laser and received at an emission of 662-737 nm. The RNA-associated RBP fluorescence signal (FRET channel) was excited with a 488-nm laser and received an emission of 662-737 nm.

### Small interfering RNA (siRNA)-based *DTWD2* knockdown

To obtain *DTWD2* knockdown cell lines, we used a siRNA transfection approach. Firstly, MCF-7 cells were seeded at a density of 1×10^5^ cells per well on glass-bottom 35-mm imaging dishes overnight. When the cell confluence reached 60%-70%, 10 μl of RNATransMate and 7 μl of 20 μM siRNA were added to centrifuge tubes containing 690 μl and 693 μl of serum-free medium, respectively, followed by 1:1 mixing. The mixture was left at room temperature for 15 min, and then added to the dishes and incubated for two days. Subsequently, the cells were processed according to the COMPASS protocol described above.

For transfection efficiency experiments, cells were transfected with the FAM-siRNA and observed by fluorescence microscopy (Nikon) (**Supplementary Fig. 13**). For qRT-PCR experiments, we used a BeyoFast™ SYBR Green One-Step qRT-PCR kit (Beyotime Biotechnology) to verify that siRNA successfully interfered with the mRNA expression of the *DTWD2* gene (**Fig. 2c**).

### Colocalization imaging of glycoRNA with cell membranes and lipid rafts

To label cell membranes, we treated MCF-7 cells on imaging dishes with the Cell Plasma Membrane Staining Kit with DiD (Beyotime Biotechnology) at 37 °C for 15 min before treatment with COMPASS. To label lipid rafts, we treated MCF-7 cells on imaging dishes with 10 μg/ml of CT-B-AF594 (Thermo Fisher Scientific) on ice for 30 min before treating with COMPASS. To accomplish the imaging, a ×100 oil immersion objective was applied, and the COMPASS system was excited with a 488-nm laser. The DiD fluorescence signal and CT-B fluorescence signal were excited with a 561-nm laser.

### Cell viability assay

#### Assessing cell viability under MCR co-treatment by CCK-8

A total of 10,000 MCF-7 cells were seeded into 96-well plates. Following overnight stabilization, the cells were subjected to a 48-hour treatment with different concentrations of MCR at 37 C. Next, 10 μL of CCK-8 solution (Macklin) was added to each well, followed by a 4-hour incubation in a dark environment at 37C. Plates were detected with a 450 nm wavelength using a Synergy H1M microplate reader (BioTek).

#### Assessing cell viability under DMBA and PTX treatment by MTT

Totally 7000 MCF-10A cells were plated in 96-well plates. After stabilization overnight, the cells were treated with different concentrations of DMBA for 24 h at 37 C. After incubation, 100 μL of 0.5 mg/ml Thiazolyl Blue (Solarbio) was added to each well and incubated for 4 h at 37C in a dark environment. Plates were detected with a 490 nm wavelength after incubating with 100 μL DMSO per well for 10 min using a Synergy H1M microplate reader (BioTek).

Totally 7000 MCF-7 cells were plated in 96-well plates with increasing doses of paclitaxel for 48 h. Next, cells were incubated with 100 μL of 0.5 mg/ml Thiazolyl Blue for 4 h at 37 C in a dark environment. Subsequently, cells were incubated with 100 μL DMSO for 4 h in the dark. Plates were detected with a wavelength of 490 nm.

### Wound healing assay

MCF-10A cells were seeded in the 6-well plates at a density of 1.5×10^6^ cells per well. When the cell confluence reached around 90%, each cell monolayer was wounded using a sterile 20 μL pipette tip. After washing three times with PBS, cells were cultured in serum-free medium containing 10 μM DMBA for 24 h. Cell migration images were observed at 0, 12, and 24 h using a digital inverted microscope (Nikon). Cells without any treatment were used as a control group. The average scratch area of each group was calculated using Fiji.

### Establish the chemotherapy model

MCF-7 cells were seeded in imaging dishes at a density of 1.5×10^5^ cells per well. After washing with PBS, cells were treated with 2.5 nM paclitaxel for 48 h to establish a chemotherapy model, followed by glycoRNA analysis using COMPASS. Notably, Cells without paclitaxel treatment were used as a control group. Notably, metabolic markers were co-incubated with cells for 24 h.

### Establish the malignancy transformation model

MCF-10A cells were seeded in imaging dishes at a density of 1.5×10^5^ cells. MCF-7 cells were seeded in imaging dishes at a density of 1.5×10^5^ cells per well. After washing with PBS, cells were treated with 10 μM DMBA for 24 h to establish a malignancy transformation model, followed by glycoRNA analysis using COMPASS. Cells without DMBA treatment were used as a control group.

### Culture of human cerebral organoid

The hESC line H9 (WiCell agreement no. 16-W0060) was used in this study. A series of manipulations was performed according to previous studies^44^. Briefly, iPSC lines were maintained on the culture plates coated with vitronectin (ThermoFisher Scientific) under feeder-free conditions. After being cultured in E8 medium (ThermoFisher Scientific) for 5 to 7 days, hPSCs were dissociated using EDTA (Lonza) at 37°C for 1-2 min and then plated in a 6-well dish at a density of 1×10^5^ cells per well. To obtain embryoid bodies (EBs) for neural differentiation, iPSCs were detached using dispase (ThermoFisher Scientific). Subsequently, these EBs were cultured in a neural induction medium comprising N2 supplement (Thermo Fisher Scientific), nonessential amino acids (MEM-NEAA, Thermo Fisher Scientific), and DMEM/F12 (ThermoFisher Scientific) for 7 days. On day 7, the EBs were resuspended in Matrigel (Corning), which was cold-pipetted into 3 mm dimples on a UV-sterilized Parafilm sheet for 30 min. These droplets solidified at 37°C and were then taken off the Parafilm to be grown in a differentiation medium, which was renewed every 5 days.

### COMPASS-assisted glycoRNA imaging in human cerebral organoid

Human cerebral organoids were incubated with Ac_4_ManNAz (50 μM) and 5-EU (250 μM) in a neural induction medium for 48 h. After metabolic labeling, they were immobilized in 4% paraformaldehyde for a duration of 2 h and then washed three times with PBS for 10 min each time. Subsequently, organoids were immersed in a 20% sucrose solution in PBS at 4 °C overnight. Once the organoids had settled to the bottom of the tube, the soaking solution was exchanged with a 30% sucrose solution in PBS at 4 °C. Organoids were embedded in an optimal cutting temperature compound and cryosectioned at 10 μm. These tissue sections were subsequently utilized for staining procedures. In brief, organoids were incubated with 100 μΜ DBCO-AF647 for 10 min at room temperature. After washing with PBS three times, they were treated with BeyoClick™ EU RNA Synthesis Kit with Alexa Fluor 488 at 37 °C for 30 min. Finally, CLSM imaging of cells was acquired on an AX (Nikon). To accomplish the imaging, a ×40 water immersion objective was applied, and the RNA fluorescence signal (AF488 channel) was excited with a 488-nm laser and received at an emission of 500-550 nm. The Glycan fluorescence signal (AF647 channel) was excited with a 640-nm laser and received an emission of 662-737 nm. The GlycoRNA fluorescence signal (FRET channel) was excited with a 488-nm laser and received an emission of 662-737 nm.

### Animal models and COMPASS-assisted glycoRNA imaging in mouse tissues

Female BALB/c mice (4–5 weeks, 20 ± 2 g) were supplied by SPF Biotechnology Co., Ltd (Suzhou, China). Mice were housed in a specific pathogen-free environment with a temperature of 21-26 °C (daily temperature difference less than 4 °C), relative humidity of 40%-70%, and a 12-hour light-dark cycle. Feed and water were available *ad libitum*. The animal experiment protocol was approved by the Laboratory Animal Ethics Committee of Nanjing Ramda Pharmaceutical Co., Ltd (Accreditation number: IACUC-2024082901).

The experimental mice were divided into the following two groups (n = 3 per group): a control group and the model group. To establish tumor metastasis models, 4T1 cells (1×10^7^ cells) suspended in PBS were injected intravenously through the tail vein in the model group. Meanwhile, the control group was injected with PBS. As shown in **Fig. 5c**, mice were injected intraperitoneally with 100 μl of 5-EU (20 mg/ml) after tumor metastasis, and Ac_4_ManNAz (5 mg/kg) was injected intraperitoneally once daily for three consecutive days. On day 4, DBCO-AF647 (0.16 mmole/kg) was injected into the tail vein three hours before sacrificing the mice for in vivo labeling of glycan. Major organs (heart, liver, kidney, spleen, and lung) were collected and washed with saline. They were immobilized in 4% paraformaldehyde and sucrose dehydration. Cryosectioned tissues were permeabilized with 0.1% Triton for 30 min and washed three times with PBS. Then, they were treated with BeyoClick™ EU RNA Synthesis Kit with Alexa Fluor 488 at 37 °C for 30 min. CLSM imaging of cells was acquired on an AX (Nikon). To accomplish the imaging, a ×20 water immersion objective was applied, and the RNA fluorescence signal (AF488 channel) was excited with a 488-nm laser and received at an emission of 500-550 nm. The Glycan fluorescence signal (AF647 channel) was excited with a 640-nm laser and received an emission of 662-737 nm. The GlycoRNA fluorescence signal (FRET channel) was excited with a 488-nm laser and received an emission of 662-737 nm.

### Data analysis

All experiments were replicated at least three times. Image analysis was processed with Fiji. The results of each test are displayed as the mean ± s.d. A two-tailed unpaired Student’s t-test and unpaired t-test were conducted to compare two independent groups. The t-test indicated no statistical significance at the thresholds of P < 0.05 (*), P < 0.01 (**), P < 0.001 (***), and P < 0.0001 (****). Statistical analyses and graphical representations were carried out using OriginPro 2024b or GraphPad Prism 10.

