## Supplementary Information for "Sequence-Independent In Situ Imaging of GlycoRNA in Living Cells and Tissues based on COMPASS"

**Table of contents**

### **Supplementary Fig. 1 |** The chemical structure of a synthetic Y5 glycoRNA for dynamic simulation.

**Supplementary Fig. 2 |** The chemical structure of a physiologically relevant Y5 glycoRNA model for dynamic simulation.

**Supplementary Fig. 3 |** Toxicity testing of MCRs.

**Supplementary Fig. 4 |** Analysis of Ac_4_ManNAz incorporation into cellular sialic acid.

**Supplementary Fig. 5 |** Analysis of 5-EU incorporation into RNA.

**Supplementary Fig. 6 |** Blotting analysis of glycoRNA extracted from MCF-7 cells.

**Supplementary Fig. 7 |** Conditional optimization of MCRs and cell incubation time.

**Supplementary Fig. 8 |** Conditional optimization of DBCO-AF647 concentration in COMPASS.

**Supplementary Fig. 9 |** Conditional optimization of DBCO-AF647 incubation time in COMPASS.

**Supplementary Fig. 10 |** Conditional optimization of N_3_-AF488 incubation concentration in COMPASS.

**Supplementary Fig. 11 |** Conditional optimization of N_3_-AF488 incubation time in COMPASS.

**Supplementary Fig. 12 |** Cross-reaction validation between two-step click chemistry.

**Supplementary Fig. 13 |** Fluorescence images of MCF-7 cells transfected with FAM-siRNA.

**Supplementary Fig. 14 |** CCK-8 assessment of MCF-7 cells treated with siRNA and MCRs.

**Supplementary Fig. 15 |** 3D visualization of the spatial distributions of glycoRNAs in MCF-7 cells.

**Supplementary Fig. 16 |** The 3D visualization of glycoRNAs with COMPASS and plasma membrane with DiD in MCF-7 cells.

**Supplementary Fig. 17 |** The 3D visualization of glycoRNAs with COMPASS and lipid rafts with CT-B in MCF-7 cells.

**Supplementary Fig. 18 |** CLSM images of RBPs on the surface of untreated, proteinase K-treated, and trypsin-treated MCF-7 cells.

**Supplementary Fig. 19 |** CLSM images of MCF-7 cells treated with COMPASS and a COMPASS-like procedure using immunofluorescence to label cell surface RBPs.

**Supplementary Fig. 20 |** Generality for COMPASS to detect glycoRNAs on the cell surface of PC-12 cells.

**Supplementary Fig. 21 |** Quantification analysis of RNA and glycan signaling in MCF-7 and MCF-10A cells.

**Supplementary Fig. 22 |** MTT assessment of MCF-10A cells treated with DMBA in a dose-dependent manner.

**Supplementary Fig. 23 |** Wound healing assay of MCF-10A cells treated with DMBA.

**Supplementary Fig. 24 |** RT-qPCR assay for *p53* and *EGFR* expression in MCF-10A cells treated with different concentrations of DMBA.

**Supplementary Fig. 25 |** Quantification of the average fluorescence intensity of RNA and glycan signaling after treatment of MCF-10A with different concentrations of DMBA**.**

**Supplementary Fig. 26 |** MTT assessment of MCF-7 cells treated with paclitaxel in a dose-dependent manner.

**Supplementary Fig. 27 |** RT-qPCR assay for *p53* and *EGFR* expression in MCF-7 cells treated with different concentrations of paclitaxel.

**Supplementary Fig. 28 |** Quantification of the average fluorescence intensity of RNA and glycan signaling after treatment of MCF-7 with different concentrations of paclitaxel.

**Supplementary Fig. 29 |** Photographs of human cerebral cortex organoid**.**

**Supplementary Fig. 30 |** CLSM imaging of 4T1 cell surface glycoRNAs with COMPASS.

**Supplementary Fig. 31 |** HPLC analysis of glycoRNA from various mouse tissues.

**Supplementary Fig. 32 |** Hematoxylin & eosin (H&E) staining of tissues from the major organs of control and model mice.

**Supplementary Fig. 33 |** CLSM images of glycoRNAs in the heart section of different treated mice by COMPASS.

**Supplementary Fig. 34 |** CLSM images of glycoRNAs in the liver section of different treated mice by COMPASS.

**Supplementary Fig. 35 |** CLSM images of glycoRNAs in the spleen section of different treated mice by COMPASS.

**Supplementary Fig. 36 |** CLSM images of glycoRNAs in the kidney section of different treated mice by COMPASS.

**Supplementary Table 1.** The comparison between COMPASS and existing glycoRNA imaging methods

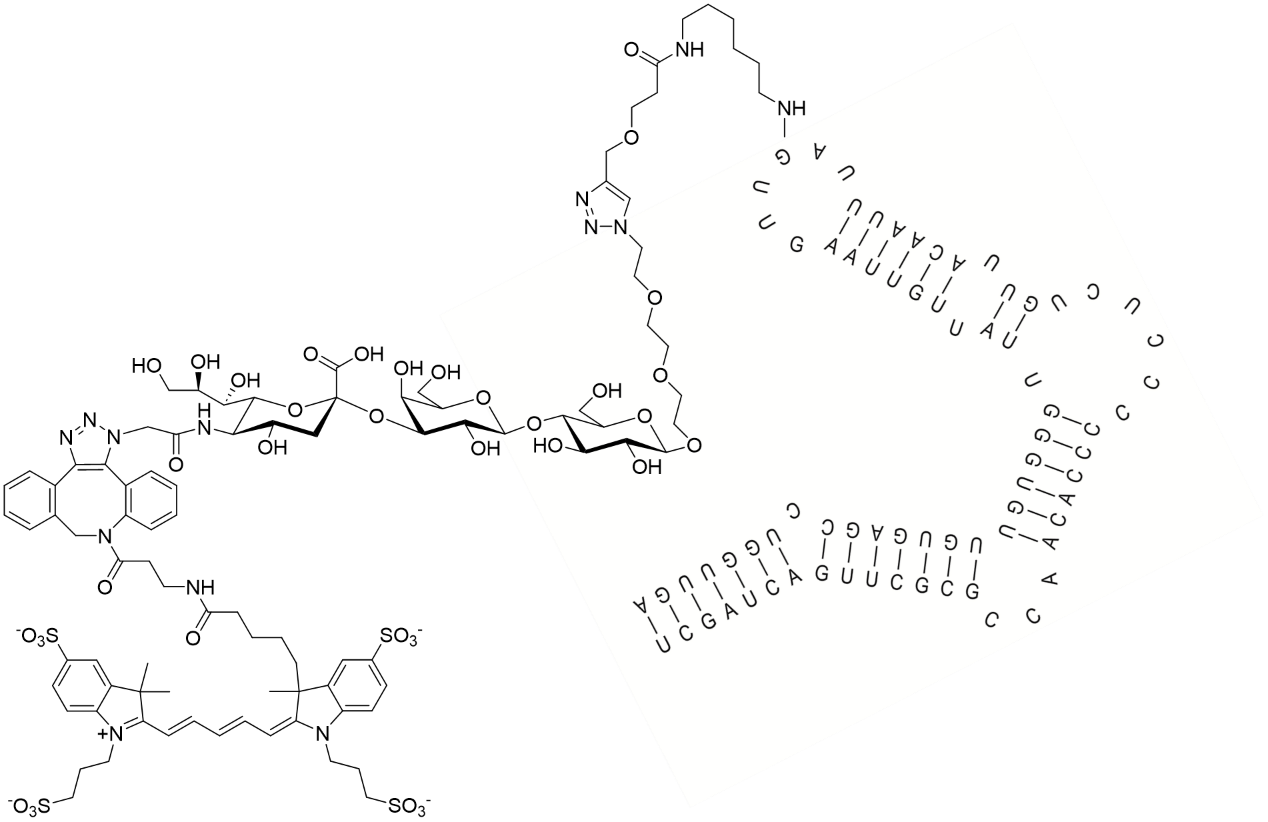

**Supplementary Fig. 1 |** **The chemical structure of a synthetic Y5 glycoRNA for dynamic simulation.**

Given the unclear nature of glycoRNA structures, particularly the actual linker between RNAs and glycans, we employed a chemical scaffold from a previous study focused on the chemical synthesis of glycoRNAs^1^. While this scaffold does not represent the natural glycoRNA structure, its impact on analyzing the distances between functional groups is minimal, as current evidence suggests that the linkage between RNA and glycan predominantly involves modified RNA connected to small organic molecules^2^. According to the reported chemical scaffold, Ac_4_ManNAz was metabolically labeled to introduce an N_3_ at the sialic acid terminus, which was then conjugated with DBCO-AF647 via click chemistry. Structural diagrams were generated using ChemDraw.

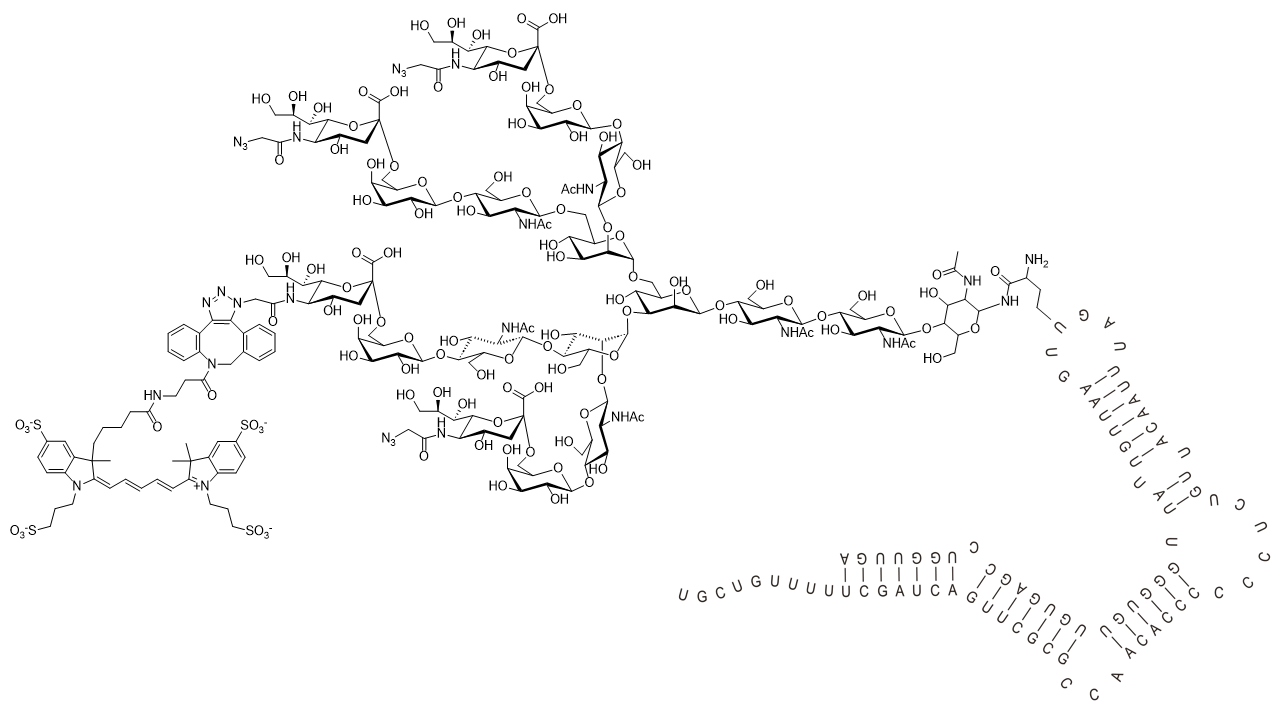

**Supplementary Fig. 2 |** **The chemical structure of** **a physiologically relevant Y5 glycoRNA model for dynamic simulation.**

Recent studies have identified acp^3^U as the linker connecting the glycan to RNA in glycoRNAs. To construct a physiologically relevant model, we performed assembly simulations using the known Y5 RNA sequence and the most abundant glycan structure. Since the precise attachment site of acp^3^U within the RNA remains undetermined, we selected a central position in the RNA sequence for linkage. Based on the reported chemical scaffolding, Ac_4_ManNAz was introduced with N_3_ at the sialic acid terminus via metabolic labeling, and then we selected an azide by click chemistry to conjugate with DBCO-AF647 for computational simulations. Structural diagrams were generated using ChemDraw.

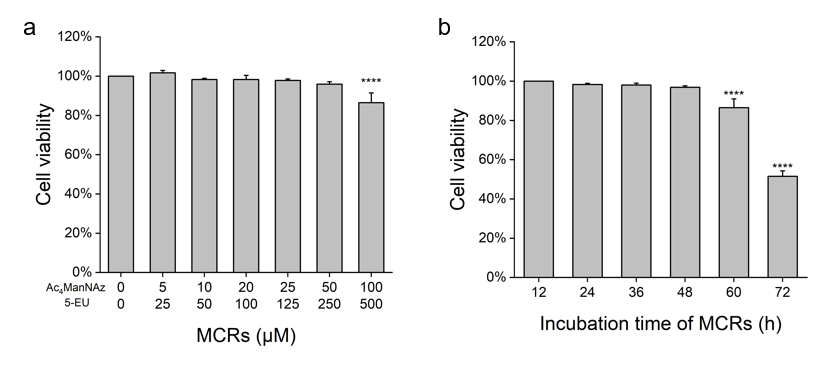

**Supplementary Fig. 3 |** **Toxicity testing of MCRs.**

**(a)** CCK-8 assessment of MCF-7 cells treated with MCRs in a dose-dependent manner. MCF-7 cells were subjected to a 48-hour treatment with different concentrations of MCRs at 37 ℃. Next, 10 μL of CCK-8 solution (Macklin) was added to each well, followed by a 4-hour incubation in a dark environment at 37 ℃. Subsequently, the plates were detected at 450 nm using a Synergy H1M microplate reader. **(b)** CCK-8 assessment of MCF-7 cells treated with MCRs in a time-dependent manner**.** MCF-7 cells were subjected to different incubation time of MCRs, including 12 h, 24 h, 36 h, 48 h, 60 h, and 72 h. Next, 10 μL of CCK-8 solution was added to each well, followed by a 4-hour incubation in a dark environment at 37 ℃. Subsequently, the plates were detected at 450 nm using a Synergy H1M microplate reader. The statistical significance is determined by unpaired two-tailed Student’s t-test as (NS) not significant, (*) P < 0.05, (**) P < 0.01, (***) P < 0.001, and (****) P < 0.0001. Data are shown as mean ± s.d.

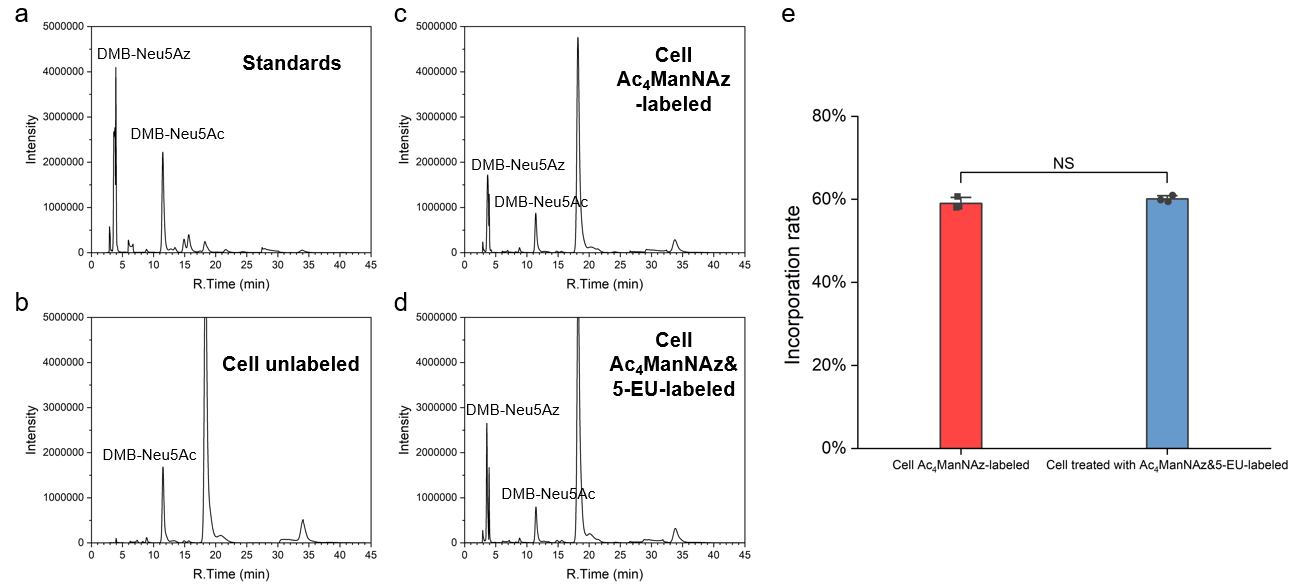

**Supplementary Fig. 4 |** **Analysis of** **Ac_4_ManNAz incorporation into cellular sialic acid.**

**(a)** HPLC separation of DMB-labeled sialic acid derivative standards from equimolar mixtures: Neu5Az and Neu5Ac. Standards were treated with DMB, and quantified by HPLC using a fluorescence detector (λ_ex_=373 nm, λ_em_=448 nm). **(b)** HPLC separation of DMB-labeled sialic acid derivative from MCF-7 cells. **(c)** HPLC separation of DMB-labeled sialic acid derivative from MCF-7 cells treated with Ac_4_ManNAz. **(d)** HPLC separation of DMB-labeled sialic acid derivative from MCF-7 cells treated with Ac_4_ManNAz and 5-EU. **(e)** Incorporation rate of azide-labeled sialic acid after treatment of cells with unnatural sugars. The statistical significance is determined by an unpaired t-test as (NS) not significant, (*) P < 0.05, (**) P < 0.01, (***) P < 0.001, and (****) P < 0.0001. Data are shown as mean ± s.d.

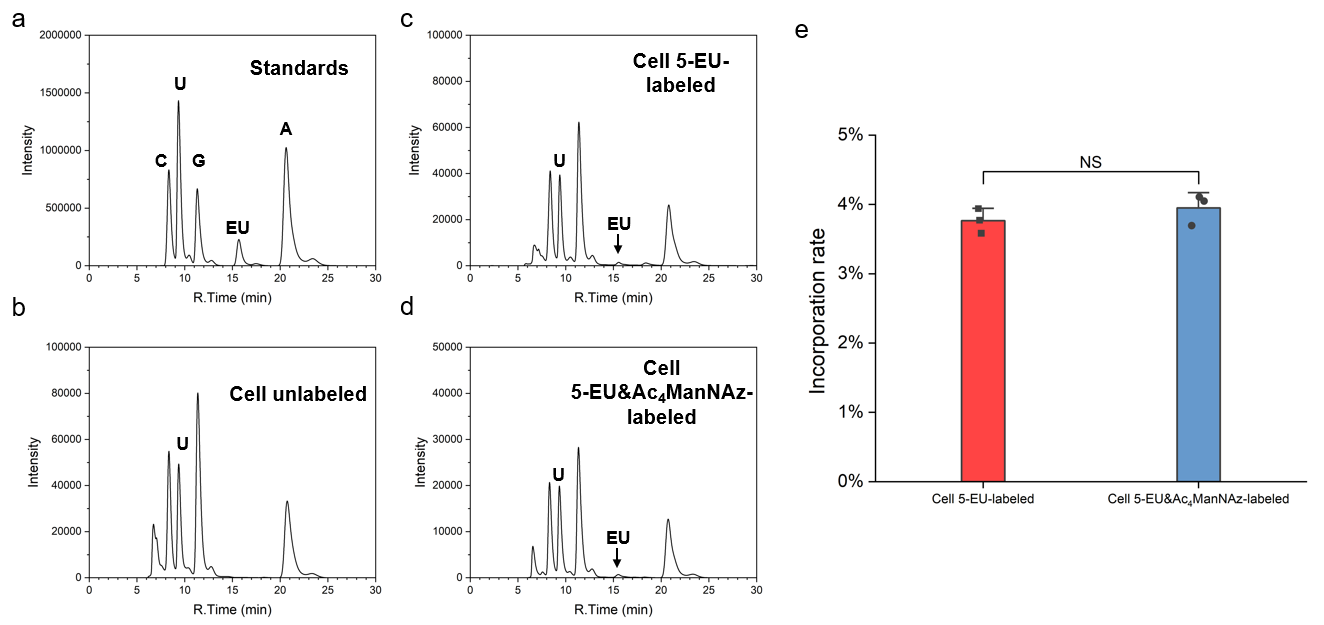

**Supplementary Fig. 5 |** **Analysis of 5-EU incorporation into RNA.**

**(a)** HPLC separation of an equimolar mixture of pure cytidine, uridine, guanosine, 5- EU, and adenosine at 260 nm. **(b)** HPLC separation of total RNA from MCF-7 cells. **(c)** HPLC separation of total RNA from MCF-7 cells treated with 5-EU. **(d)** HPLC separation of total RNA from MCF-7 cells treated with 5-EU and Ac_4_ManNAz. **(e)** Incorporation rate of alkyne-labeled RNA after treatment of cells with 5-EU. The statistical significance is determined by an unpaired t-test as (NS) not significant, (*) P < 0.05, (**) P < 0.01, (***) P < 0.001, and (****) P < 0.0001. Data are shown as mean ± s.d.

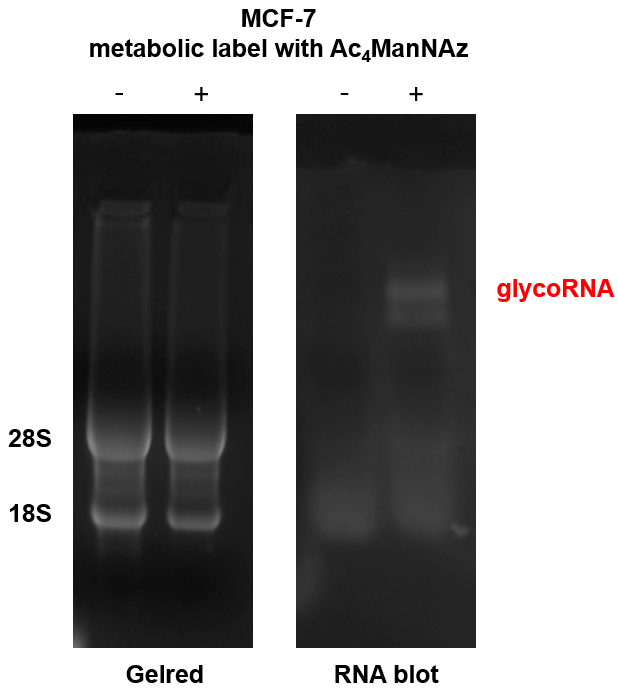

**Supplementary Fig. 6 | Blotting analysis of glycoRNA extracted from MCF-7 cells.**

Blotting of total RNA from MCF-7 cells after metabolic labeling with Ac_4_ManNAz, or MCF-7 cells without metabolic labeling.

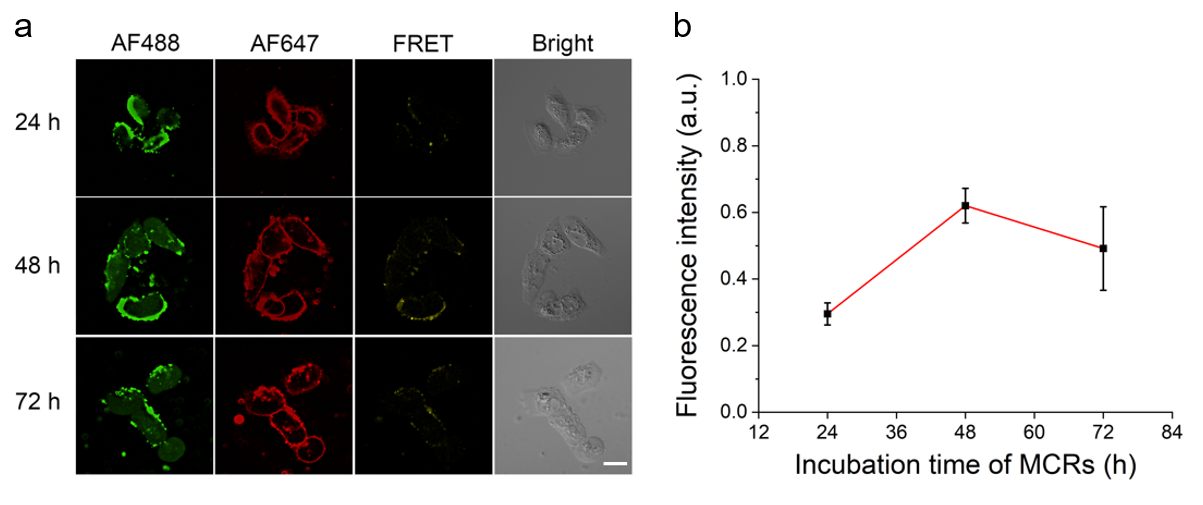

**Supplementary Fig. 7 |** **Conditional optimization of MCRs and cell incubation time.**

**(a)** CLSM images of MCF-7 cells incubated with MCRs for different time, including 24 h, 48 h, and 72 h. Cells were subsequently treated with SPAAC reaction. Incubation of MCF-7 cells at 37 °C for another 30 min for CuAAC reaction. Scale bar, 20 µm. **(b)** Statistical FRET average fluorescence intensities from **(a)** with Nikon AX treated with Fiji, n = 5 frames. Data are shown as mean ± s.d.

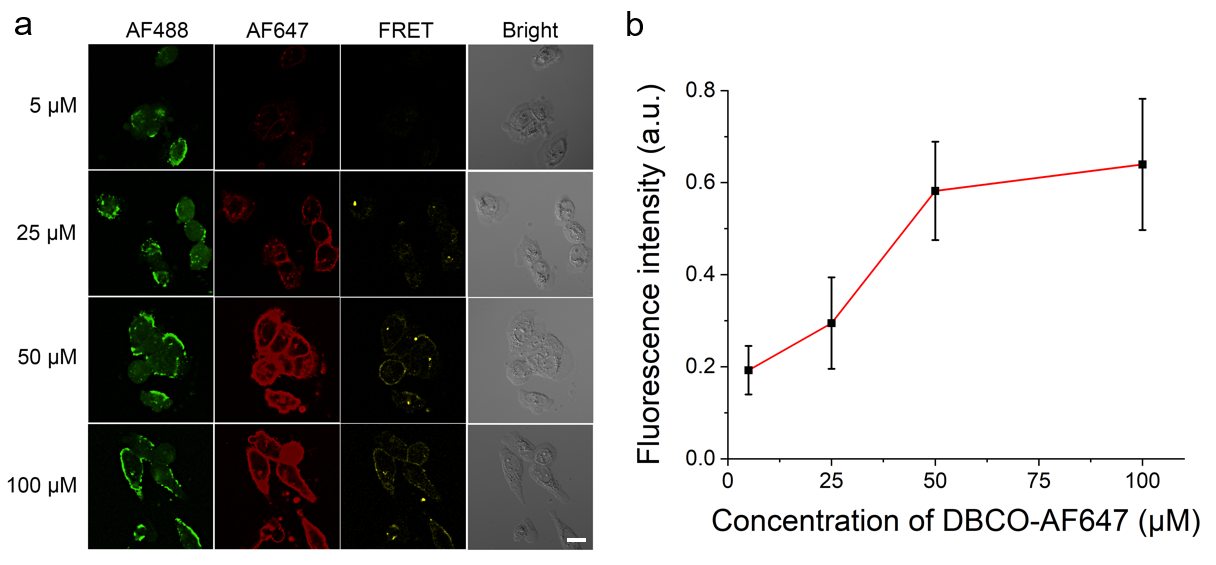

**Supplementary Fig. 8 |** **Conditional optimization of DBCO-AF647 concentration in COMPASS.**

**(a)** MCF-7 cells were incubated with 50 μM Ac_4_ManNAz and 250 μM 5-EU for 48 h in a cell culture incubator. Cells were subsequently treated with different concentrations of DBCO-AF647, containing 5 μM, 25 μM, 50 μM, and 100 μM, on ice for 10 min. Incubation of MCF-7 cells at 37 °C for another 30 min for CuAAC reaction. Scale bar, 20 μm. **(b)** Statistical FRET average fluorescence intensities from **(a)** with Nikon AX treated with Fiji, n = 5 frames. Data are shown as mean ± s.d.

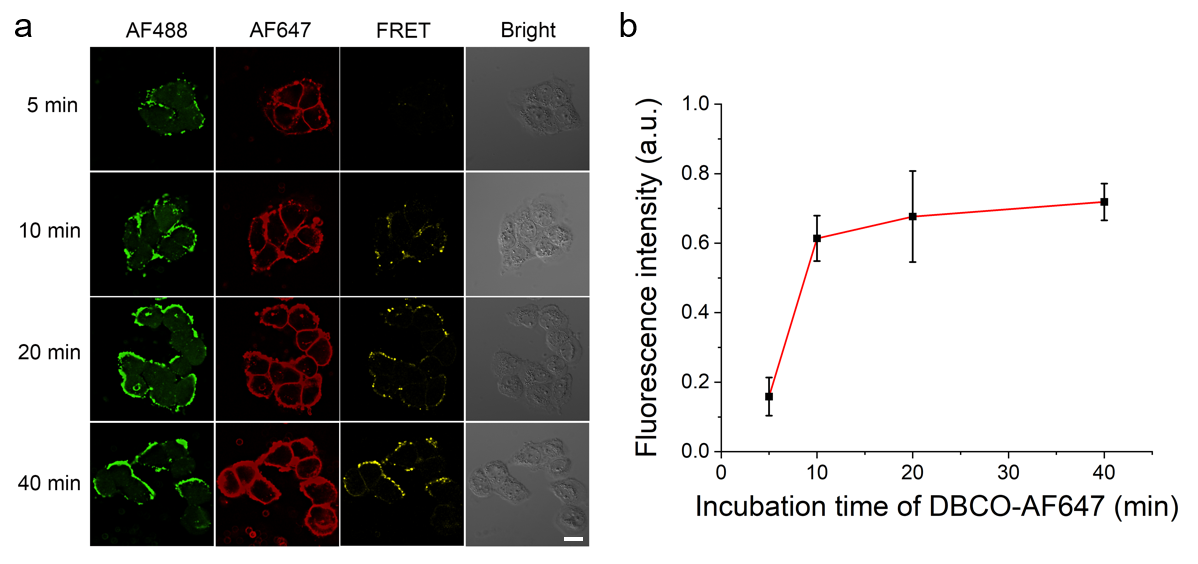

**Supplementary Fig. 9 |** **Conditional optimization of DBCO-AF647 incubation time in COMPASS.**

**(a)** MCF-7 cells were incubated with 50 μM Ac_4_ManNAz and 250 μM 5-EU for 48 h in a cell culture incubator. Cells were subsequently treated with 50 μM DBCO-AF647 on ice for different time, including 5 min, 10 min, 20 min, and 40 min. Incubation of MCF-7 cells at 37 °C for another 30 min for CuAAC reaction. Scale bar, 20 μm. **(b)** Statistical FRET average fluorescence intensities from **(a)** with Nikon AX treated with Fiji, n = 5 frames. Data are shown as mean ± s.d.

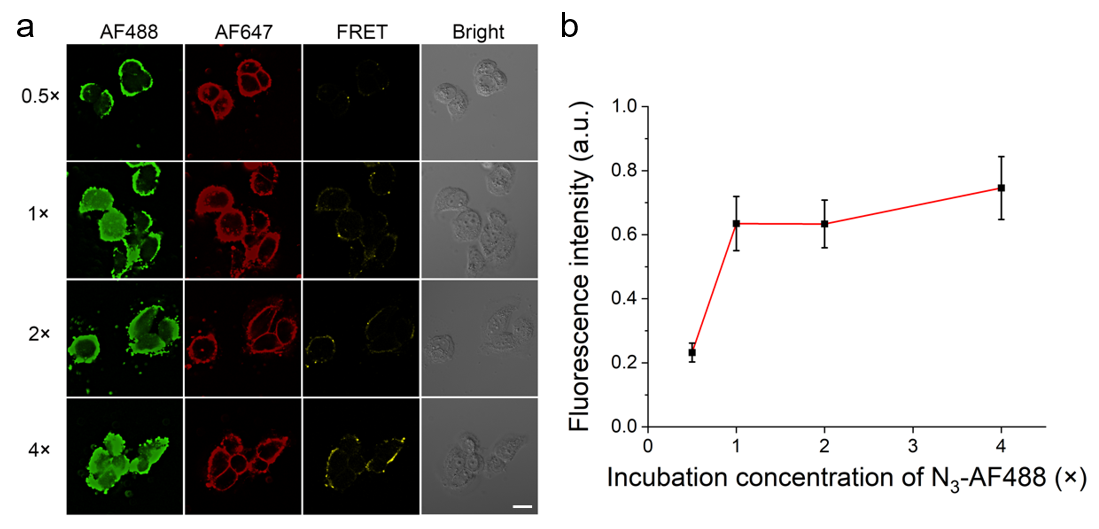

**Supplementary Fig. 10 |** **Conditional optimization of N_3_-AF488 incubation concentration in COMPASS.**

**(a)** MCF-7 cells were incubated with 50 μM Ac_4_ManNAz and 250 μM 5-EU for 48 h in a cell culture incubator. Cells were subsequently treated with SPAAC reaction. Then, cells were treated with 0.5-4× N_3_-AF488 at 37 °C for 30 min. Scale bar, 20 μm. **(b)** Statistical FRET average fluorescence intensities from **(a)** with Nikon AX treated with Fiji, n = 5 frames. Data are shown as mean ± s.d.

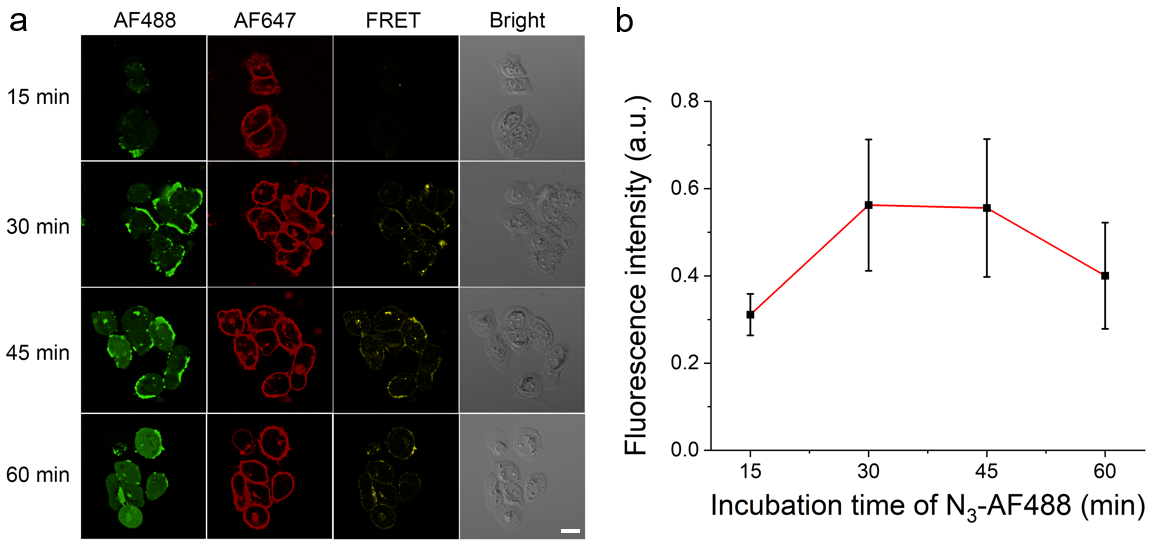

**Supplementary Fig. 11 |** **Conditional optimization of N_3_-AF488 incubation time in COMPASS.**

**(a)** MCF-7 cells were incubated with 50 μM Ac_4_ManNAz and 250 μM 5-EU for 48 h in a cell culture incubator. Cells were subsequently treated with SPAAC reaction. Then, cells were treated with 1× N_3_-AF488 at 37 °C for different time, including 15 min, 30 min, 45 min, and 60 min. Scale bar, 20 μm. **(b)** Statistical FRET average fluorescence intensities from **(a)** with Nikon AX treated with Fiji, n = 5 frames. Data are shown as mean ± s.d.

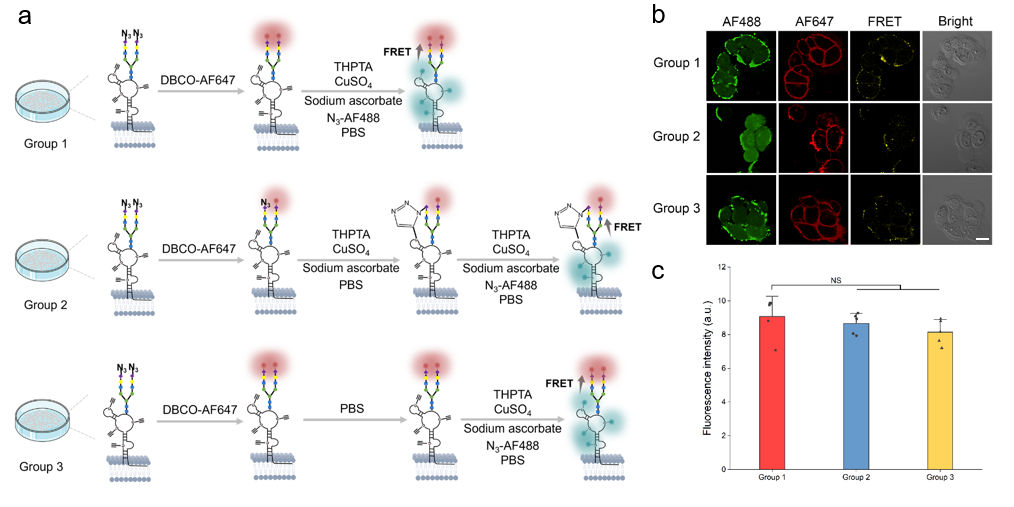

**Supplementary Fig. 12 |** **Cross-reaction validation between two-step click chemistry.**

**(a)** Schematic of a cross-reaction validation experiment between two-step click chemistry. A series of controlled experiments involving three parallel groups were designed: (1) the standard COMPASS protocol; (2) cells labeled with DBCO-AF647, followed by the addition of CuAAC reaction components (excluding N_3_-AF488), and then subsequent introduction of N_3_-AF488; and (3) cells labeled with DBCO-AF647, followed by buffer incubation (as a control for the CuAAC reaction components), and then introduction of N_3_-AF488 along with the CuAAC components. **(b)** CLSM images of MCF-7 cells treated according to the experimental steps in **(a)**. Scale bar, 20 µm. **(c)** Statistical AF488 fluorescence intensities from **(b)** with Fiji. The statistical significance is determined by unpaired two-tailed Student’s t-test as (NS) not significant, (*) P < 0.05, (**) P < 0.01, (***) P < 0.001, and (****) P < 0.0001. Data are shown as mean ± s.d.

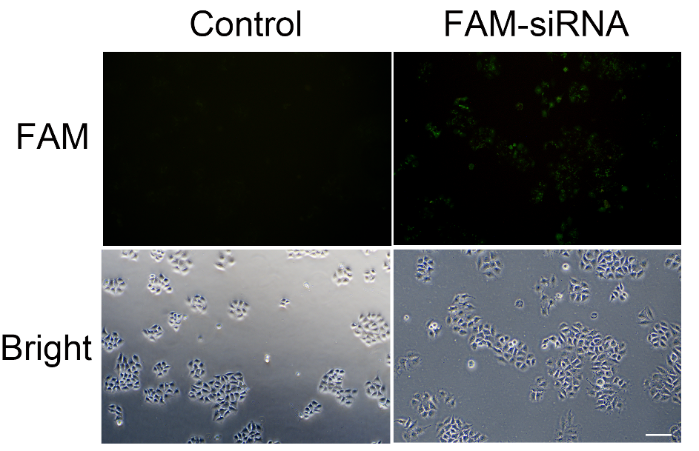

**Supplementary Fig. 13 | Fluorescence images of MCF-7 cells transfected with FAM-siRNA.**

Totally 5 nM FAM-siRNA was transfected into MCF-7 cells, and good transfection efficiency was found under these conditions. Scale bar, 100 µm.

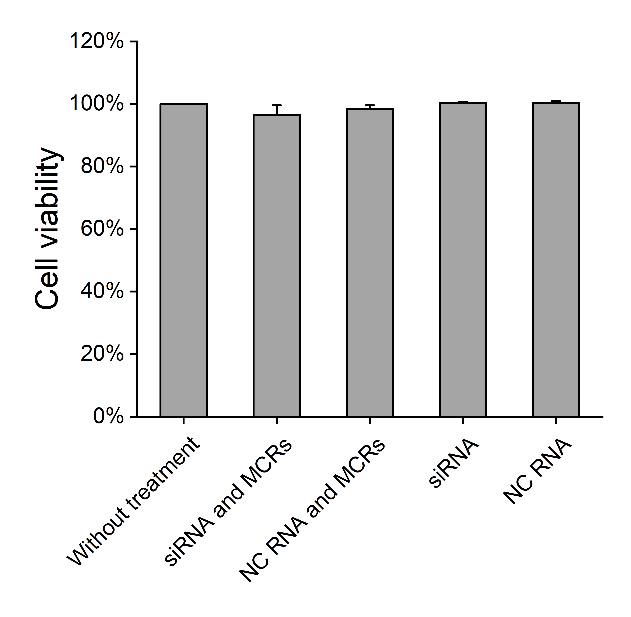

**Supplementary Fig. 14 | CCK-8 assessment of MCF-7 cells treated with siRNA and MCRs.**

MCF-7 cells were subjected to different reagents, including siRNA and MCRs, negative control (NC) RNA and MCRs, siRNA, and NC RNA for 48 h. Next, 10 μL of CCK-8 solution (Macklin) was added to each well, followed by a 4-hour incubation in a dark environment at 37 ℃. Subsequently, the plates were detected at 450 nm using a Synergy H1M microplate reader.

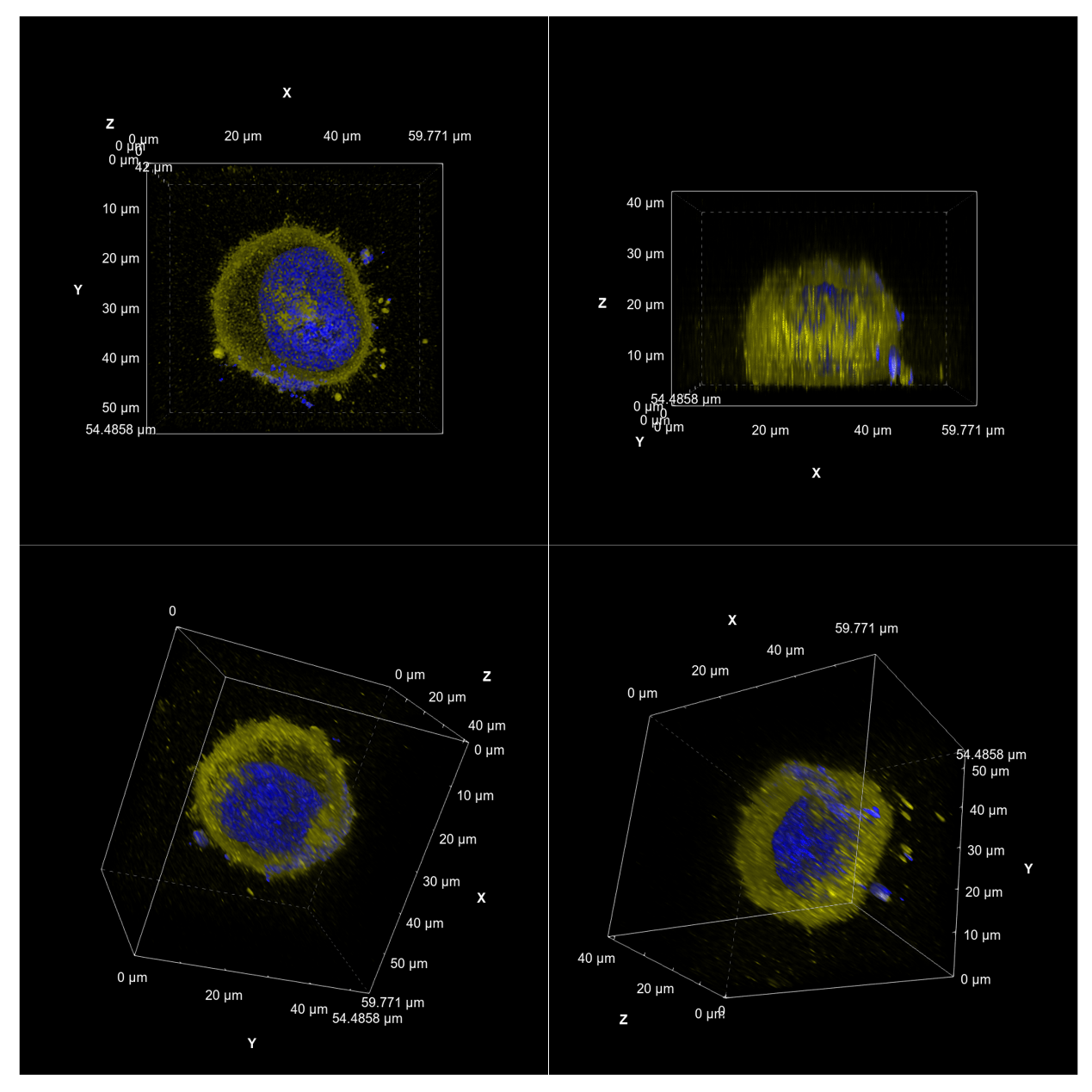

**Supplementary Fig. 15 | 3D visualization of the spatial distributions of glycoRNAs in MCF-7 cells.**

These cells were metabolically labeled by Ac_4_ManNAz and 5-EU for 48 h and then treated with Hoechst, followed by COMPASS-assisted fluorescent imaging. The FRET signals derived from COMPASS are presented in yellow, and the nucleus stained by Hoechst is indicated in blue.

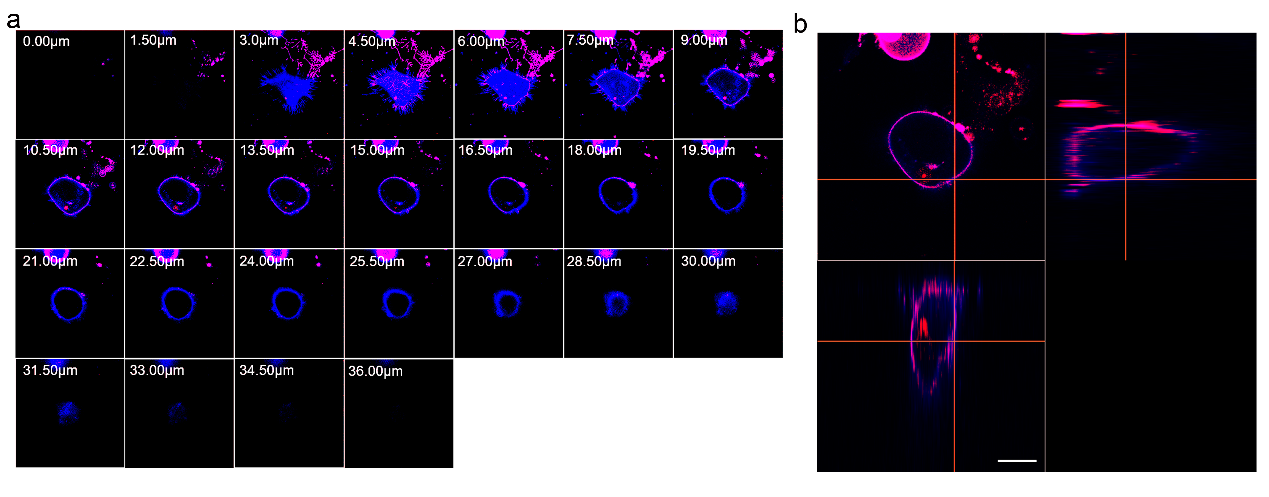

**Supplementary Fig. 16 |** **The 3D visualization of glycoRNAs with COMPASS and** **plasma membrane with DiD in MCF-7 cells.**

Z-stack images were collected with the staining of glycoRNAs by COMPASS (violet) and the plasma membrane by DiD (blue). The images were shown in z-slices format (A) and orthographic projection (B). Scale bar, 2 μm.

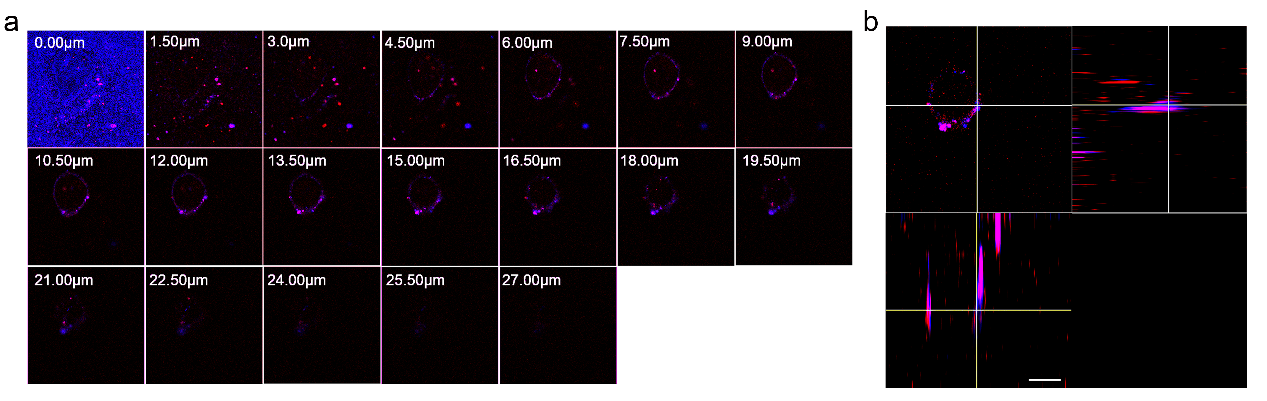

**Supplementary Fig. 17 |** **The 3D visualization of glycoRNAs with COMPASS and lipid rafts with CT-B in MCF-7 cells.**

Z-stack images were collected with the staining of glycoRNAs by COMPASS (violet) and lipid rafts by CT-B (blue). The images were shown in z-slices format (A) and orthographic projection (B). Scale bar, 2 μm.

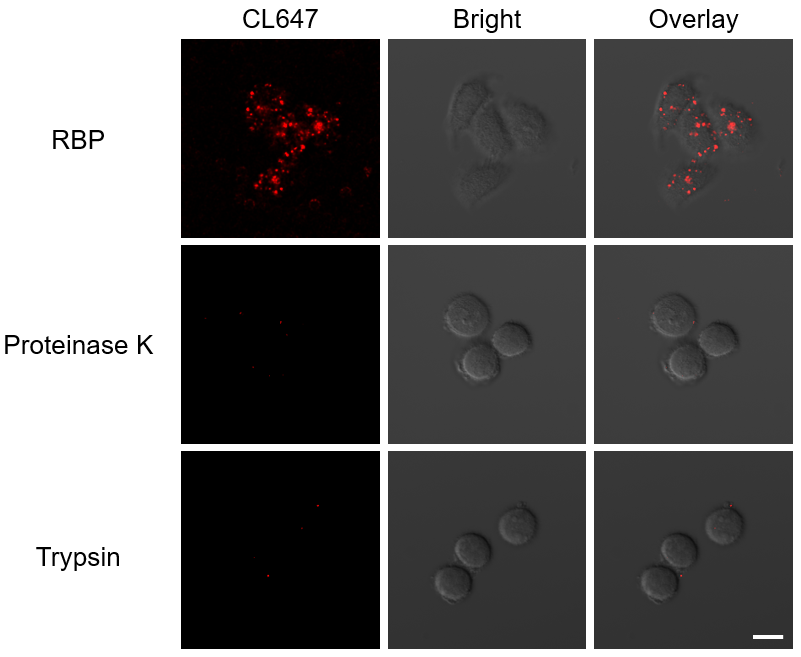

**Supplementary Fig. 18 |** **CLSM images of RBPs on the surface of untreated, proteinase K-treated, and trypsin-treated MCF-7 cells.**

For RBP imaging, MCF-7 cells were incubated with 2.5 mg/mL anti-hnRNP U antibody (Proteintech) for 45 min on ice, followed by incubation with CoraLite647-conjugated F(ab')2 Fragment Goat Anti-Rabbit IgG (H+L) antibody (Proteintech) for 30 min. Finally, cells were imaged using a confocal laser scanning microscope. In proteinase K and trypsin treatment experiments, MCF-7 cells were incubated with trypsin or 0.02 mg/mL proteinase K at 37 °C for 30 s and then analyzed by 2.5 mg/mL anti-hnRNP U antibody and CoraLite647-conjugated F(ab')2 Fragment Goat Anti-Rabbit IgG (H+L) antibody. Finally, cells were imaged using a confocal laser scanning microscope. Scale bar, 20 µm.

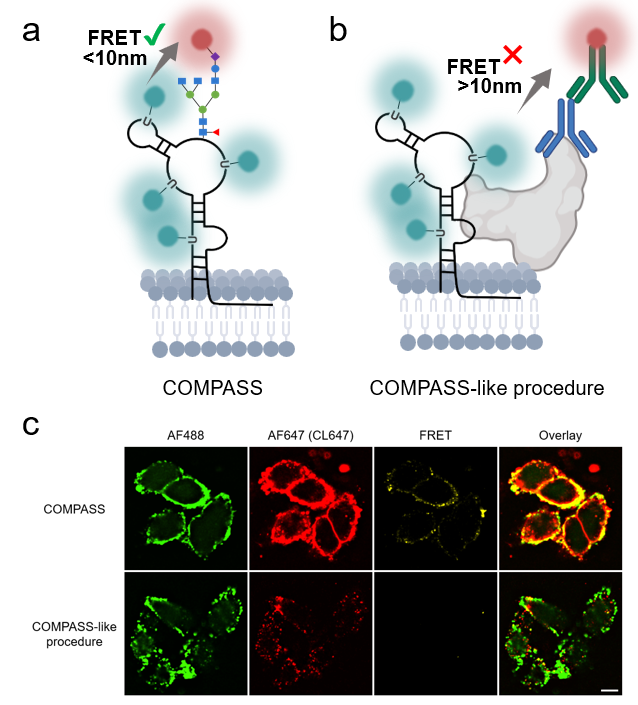

**Supplementary Fig. 19 |** **CLSM images of MCF-7 cells treated with COMPASS and a COMPASS-like procedure using immunofluorescence to label cell surface RBPs.**

**(a)** Diagram of COMPASS-tagged glycoRNA. **(b)** Diagram of COMPASS-like procedure tagged RBPs and RNA. **(c)** For COMPASS-assisted glycoRNA imaging, MCF-7 cells were incubated with Ac_4_ManNAz (50 μM) and 5-EU (250 μM) in culture medium for 48 h, followed by incubation with 50 μΜ DBCO-AF647 for 10 min on ice. Then, the cells were treated with BeyoClick™ EU RNA Synthesis Kit with Alexa Fluor 488 at 37 °C for 30 min. Finally, cells were imaged using a confocal laser scanning microscope. For COMPASS-like procedure using immunofluorescence to label surface RBPs, MCF-7 cells were cultured in medium supplemented with 5-EU (250 μM) for 48 h. Then, cells were incubated with 2.5 mg/mL anti-hnRNP U antibody for 45 min on ice, followed by incubation with CoraLite647-conjugated F(ab')2 Fragment Goat Anti-Rabbit IgG (H+L) antibody for 30 min. Subsequently, cells were treated with the BeyoClick™ EU RNA Synthesis Kit (containing Alexa Fluor 488) for 30 min at 37°C. Finally, cells were imaged using a confocal laser scanning microscope following the imaging conditions as described in the COMPASS protocol. Notably, CoraLite647 (CL647) has spectral properties comparable to those of AF647, with excitation overlap with AF488 emission, thereby forming an effective FRET pair and permitting imaging under identical CLSM settings. Scale bar, 20 µm.

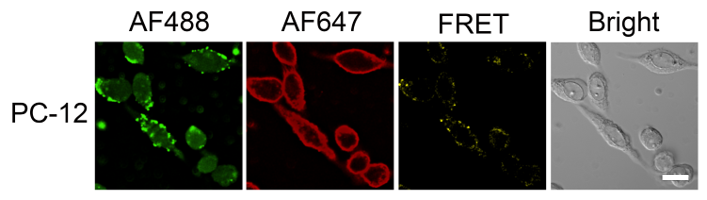

**Supplementary Fig. 20 |** **Generality for COMPASS to detect glycoRNAs on the cell surface of PC-12 cells.**

PC-12 cells were incubated with Ac_4_ManNAz (50 μM) and 5-EU (250 μM) in culture medium for 48 h. After metabolic labeling, cells were incubated with 50 μΜ DBCO-AF647 for 10 min on ice. Then, the cells were treated with BeyoClick™ EU RNA Synthesis Kit with Alexa Fluor 488 at 37 °C for 30 min. Finally, cells were imaged using a confocal laser scanning microscope. Scale bar, 20 µm.

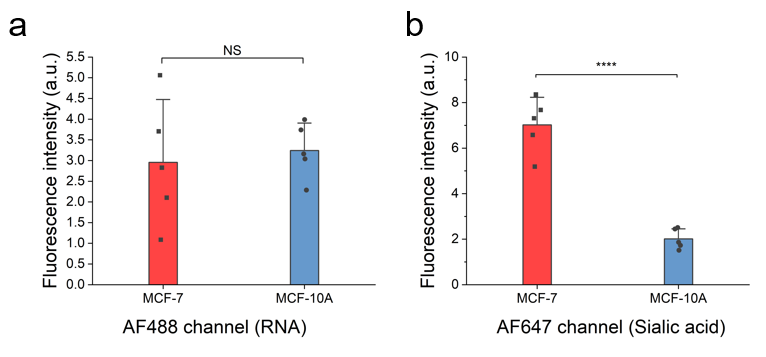

**Supplementary Fig. 21 | Quantification analysis of RNA and glycan signaling in MCF-7 and MCF-10A cells.**

**(a)** Quantification of the average fluorescence intensity of the AF488 channel in **Fig. 4a**. The AF488 channel represents the RNA signal. **(b)** Quantification of the average fluorescence intensity of the AF647 channel in Fig. 4a. The AF647 channel represents the sialic acid signal. Data is representative of three independent experiments; n = 5 frames. The statistical significance is determined by an unpaired t-test as (NS) not significant, (*) P < 0.05, (**) P < 0.01, (***) P < 0.001, and (****) P < 0.0001. Data are shown as mean ± s.d.

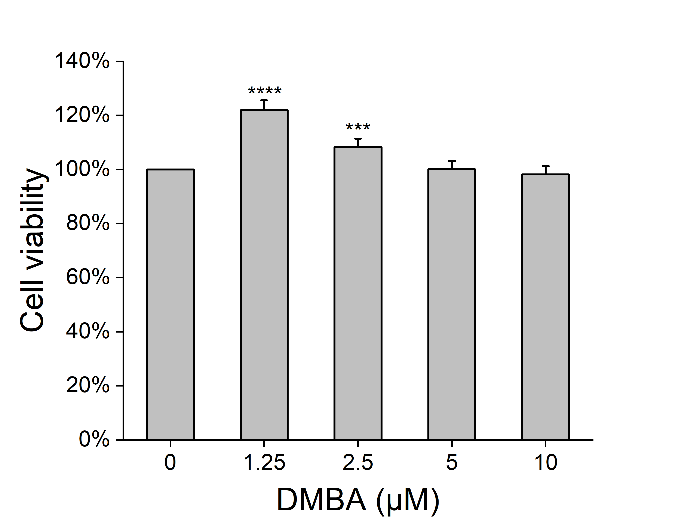

**Supplementary Fig. 22 | MTT assessment of MCF-10A cells treated with DMBA in a dose-dependent manner.**

After treating the MCF-10A cells with different concentrations (0 μM, 1.25 μM, 2.5 μM, 5 μM, 10 μM) of DMBA for 24 h, 0.5 mg/ml of Thiazolyl blue was added to each well and incubated at 37 °C for 4 h. After incubation for 10 minutes with DMSO, the plates were detected at 490 nm using a Synergy H1M microplate reader. The statistical significance is determined by unpaired two-tailed Student’s t-test as (ns) not significant, (*) P < 0.05, (**) P < 0.01, (***) P < 0.001, and (****) P < 0.0001. Data are shown as mean ± s.d.

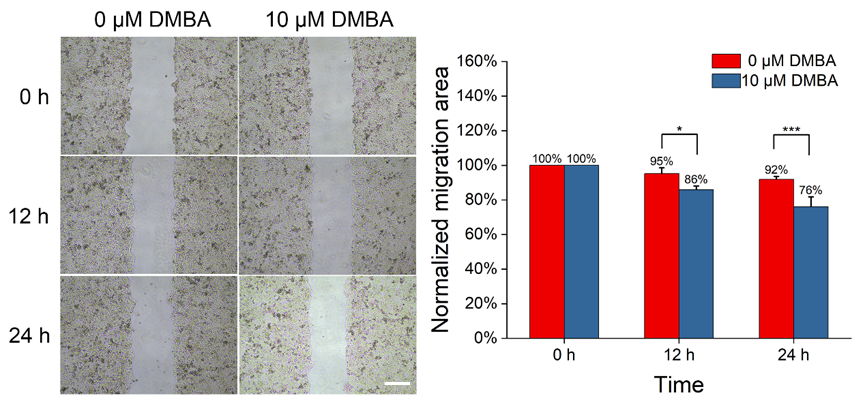

**Supplementary Fig. 23 | Wound healing assay of MCF-10A cells treated with DMBA.**

When the cell confluence reached around 100% in the 6-well plate, each cell monolayer was wounded using a sterile 20 μL pipette tip. After washing three times with PBS, cells were cultured in serum-free medium containing 10 μM DMBA for 24 h. Cell migration images were observed at 0, 12, and 24 h using a digital inverted microscope (Nikon). Cells without any treatment were used as a control group. The average scratch area of each group was calculated using Fiji. Scale bar, 100 μm. The statistical significance is determined by unpaired two-tailed Student’s t-test as (NS) not significant, (*) P < 0.05, (**) P < 0.01, (***) P < 0.001, and (****) P < 0.0001. Data are shown as mean ± s.d.

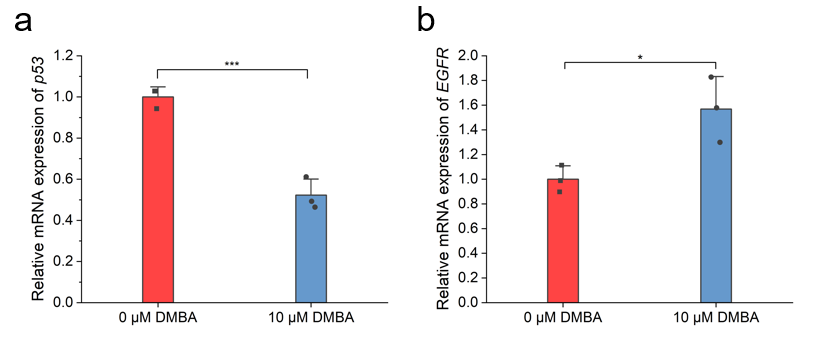

**Supplementary Fig. 24 |** **RT-qPCR assay for *p53* and *EGFR* expression in MCF-10A cells treated with different concentrations of DMBA.**

**(a)** Relative mRNA expression levels of *p53* in MCF-10A cells treated with varying concentrations of DMBA. Total RNA was extracted from MCF-10A cells after treatment with different concentrations of DMBA for 24 h. Subsequently, RT-qPCR was tested with BeyoFast™ SYBR Green One-Step qRT-PCR Kit (Beyotime) and *p53* primers. The relative mRNA expression levels were normalized to the housekeeping gene GAPDH. **(b)** Relative mRNA expression levels of *EGFR* in MCF-10A cells treated with varying concentrations of DMBA. Total RNA was extracted from MCF-10A cells after treatment with different concentrations of DMBA for 24 h. Subsequently, RT-qPCR was tested with BeyoFast™ SYBR Green One-Step qRT-PCR Kit (Beyotime) and *EGFR* primers. The relative mRNA expression levels were normalized to the housekeeping gene GAPDH. The statistical significance is determined by an unpaired t-test as (NS) not significant, (*) P < 0.05, (**) P < 0.01, (***) P < 0.001, and (****) P < 0.0001. Data are shown as mean ± s.d.

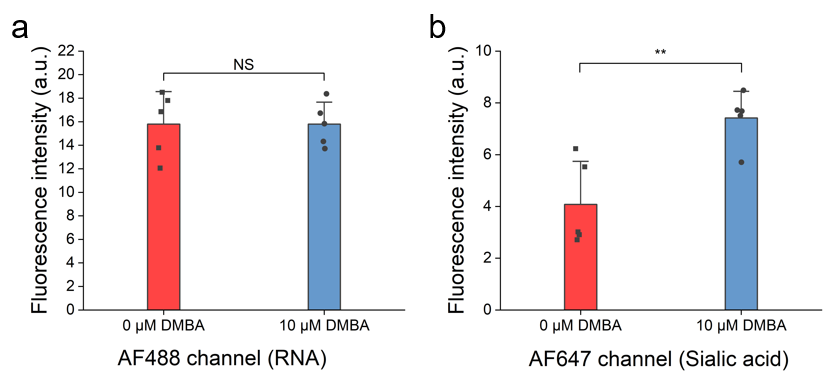
**Supplementary Fig. 25 |** **Quantification of the average fluorescence intensity of RNA and glycan signaling after treatment of MCF-10A** **with different concentrations of DMBA.**

**(a)** Quantification of the average fluorescence intensity of the AF488 channel in Fig. 4e. The AF488 channel represents the RNA signal. **(b)** Quantification of the average fluorescence intensity of the AF647 channel in Fig. 4e. The AF647 channel represents the sialic acid signal. Data is representative of three independent experiments; n = 5 frames. The statistical significance is determined by an unpaired t-test as (NS) not significant, (*) P < 0.05, (**) P < 0.01, (***) P < 0.001, and (****) P < 0.0001. Data are shown as mean ± s.d.

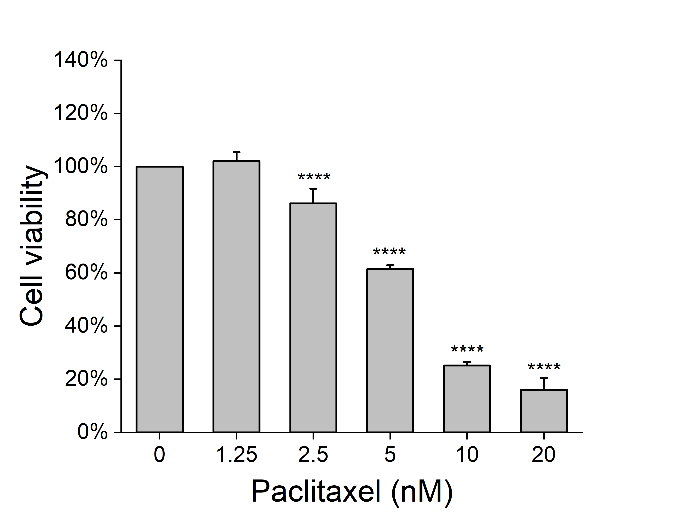

**Supplementary Fig. 26 | MTT assessment of MCF-7 cells treated with paclitaxel in a dose-dependent manner.**

After treating the MCF-7 cells with different concentrations (0 nM, 1.25 nM, 2.5 nM, 5 nM, 10 nM, 20 nM) of paclitaxel for 48 h, 0.5 mg/ml of Thiazolyl blue was added to each well and incubated at 37 °C for 4 h. After incubation for 10 minutes with DMSO, the plates were detected at 490 nm using a Synergy H1M microplate reader. The statistical significance is determined by unpaired two-tailed Student’s t-test as (NS) not significant, (*) P < 0.05, (**) P < 0.01, (***) P < 0.001, and (****) P < 0.0001. Data are shown as mean ± s.d.

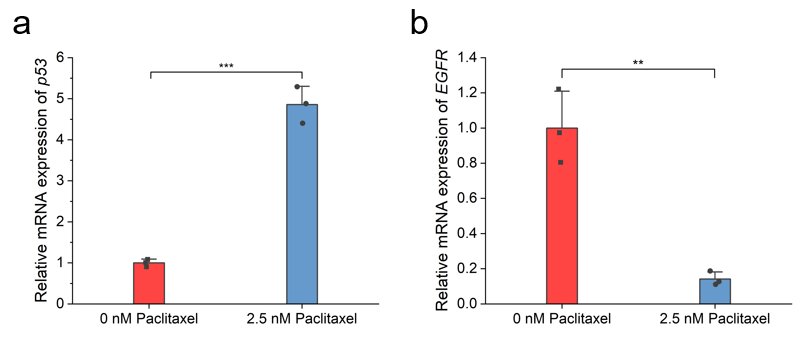

**Supplementary Fig. 27 | RT-qPCR assay for *p53* and *EGFR* expression in MCF-7 cells treated with different concentrations of paclitaxel.**

**(a)** Relative mRNA expression levels of *p53* in MCF-7 cells treated with varying concentrations of paclitaxel. Total RNA was extracted from MCF-7 cells after treatment with different concentrations of paclitaxel for 48 h. Subsequently, RT-qPCR was tested with BeyoFast™ SYBR Green One-Step qRT-PCR Kit (Beyotime) and *p53* primers. The relative mRNA expression levels were normalized to the housekeeping gene GAPDH. **(b)** Relative mRNA expression levels of *EGFR* in MCF-7 cells treated with varying concentrations of paclitaxel. Total RNA was extracted from MCF-7 cells after treatment with different concentrations of paclitaxel for 48 h. Subsequently, RT-qPCR was tested with BeyoFast™ SYBR Green One-Step qRT-PCR Kit (Beyotime) and *EGFR* primers. The relative mRNA expression levels were normalized to the housekeeping gene GAPDH. The statistical significance is determined by an unpaired t-test as (NS) not significant, (*) P < 0.05, (**) P < 0.01, (***) P < 0.001, and (****) P < 0.0001. Data are shown as mean ± s.d.

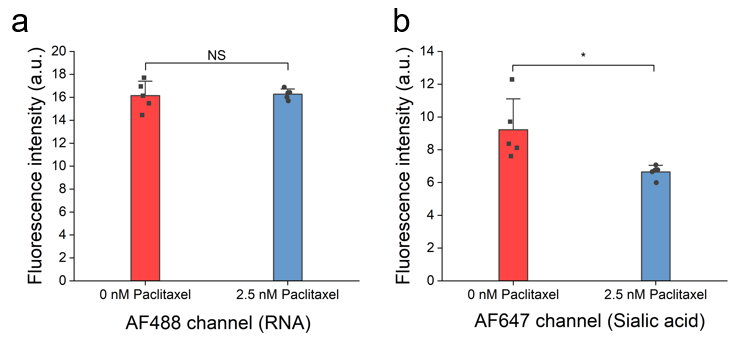

**Supplementary Fig. 28 | Quantification of the average fluorescence intensity of RNA and glycan signaling after treatment** **of MCF-7 with different concentrations of paclitaxel.**

**(a)** Quantification of the average fluorescence intensity of the AF488 channel in Fig. 4f. The AF488 channel represents the RNA signal. **(b)** Quantification of the average fluorescence intensity of the AF647 channel in Fig. 4f. The AF647 channel represents the sialic acid signal. Data is representative of three independent experiments; n = 5 frames. The statistical significance is determined by an unpaired t-test as (NS) not significant, (*) P < 0.05, (**) P < 0.01, (***) P < 0.001, and (****) P < 0.0001. Data are shown as mean ± s.d.

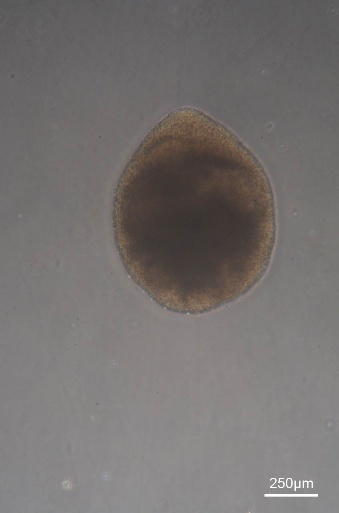

**Supplementary Fig. 29 |** **Photographs of** **human cerebral cortex organoid.**

Organoids cultured for 40 days showed a regular spherical structure with a diameter of about 750 μm. Scale bar, 250 µm.

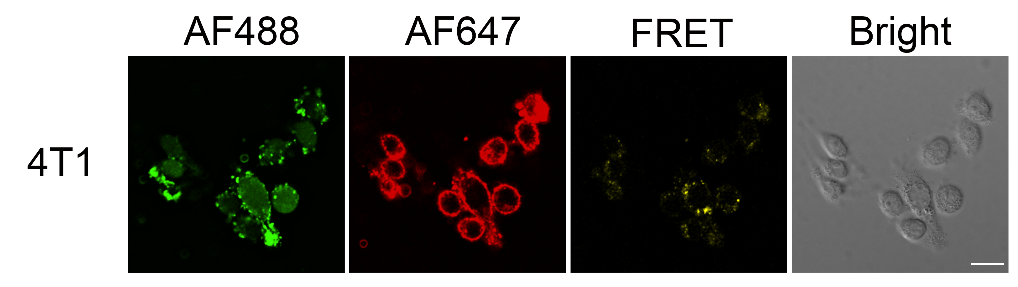

**Supplementary Fig. 30 |** **CLSM imaging of 4T1 cell surface glycoRNAs with COMPASS.**

4T1 cells were incubated with Ac_4_ManNAz (50 μM) and 5-EU (250 μM) in culture medium for 48 h. After metabolic labeling, cells were incubated with 50 μΜ DBCO-AF647 for 10 min on ice. Then, the cells were treated with BeyoClick™ EU RNA Synthesis Kit with Alexa Fluor 488 at 37 °C for 30 min. Finally, cells were imaged using a confocal laser scanning microscope. Scale bar, 20 µm.

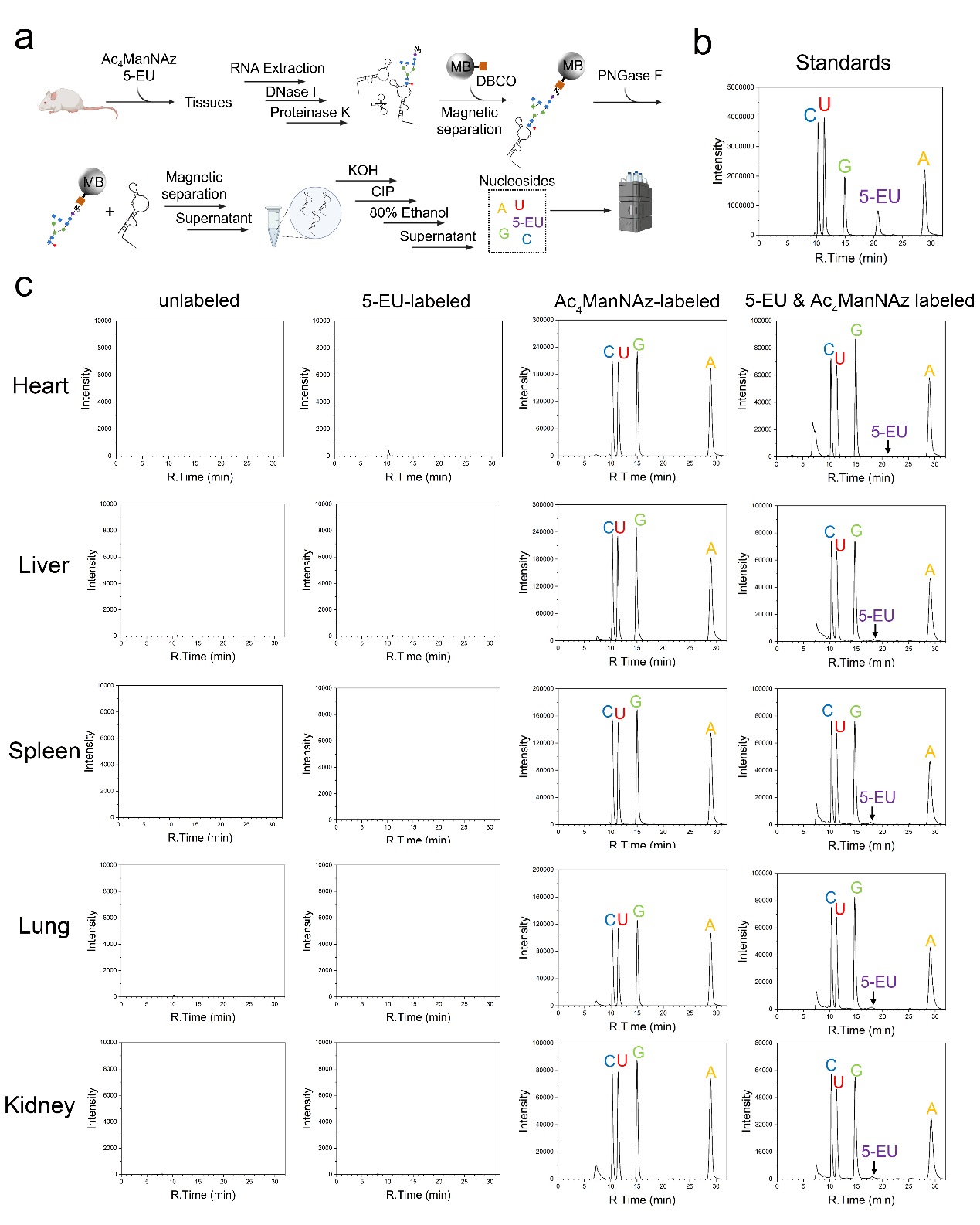

**Supplementary Fig. 31 | HPLC analysis of glycoRNA from various mouse tissues.**

**(a)** Schematic illustration of glycoRNA enrichment using DBCO-functionalized magnetic beads (MBs), followed by on-bead RNA release via PNGase F treatment, in-solution RNA hydrolysis, and HPLC analysis. **(b)** HPLC chromatograms of an equimolar mixture of nucleoside standards. **(c)** HPLC chromatograms of glycoRNA isolated from tissuess that were unlabeled, labeled with 5-EU, labeled with Ac_4_ManNAz, or co-labeled with 5-EU and Ac4ManNAz.

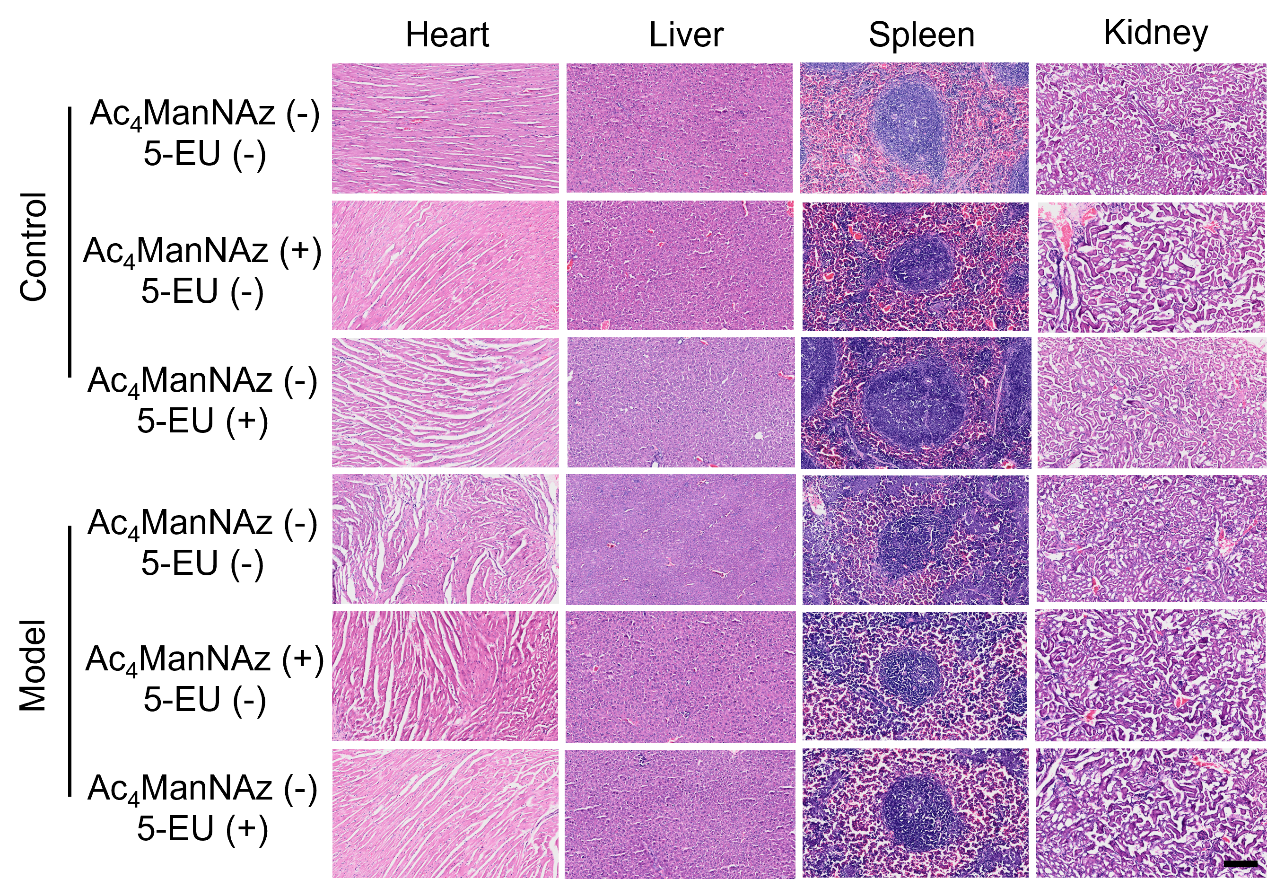

**Supplementary Fig. 32 |** **Hematoxylin & eosin (H&E) staining of tissues from the major organs of control and model mice.**

Hematoxylin stains nuclei blue-purple, while eosin highlights cytoplasmic and extracellular matrix components in pink, enabling histological assessment of tissue morphology, inflammatory infiltration, and pathological changes. Scale bar, 100 µm.

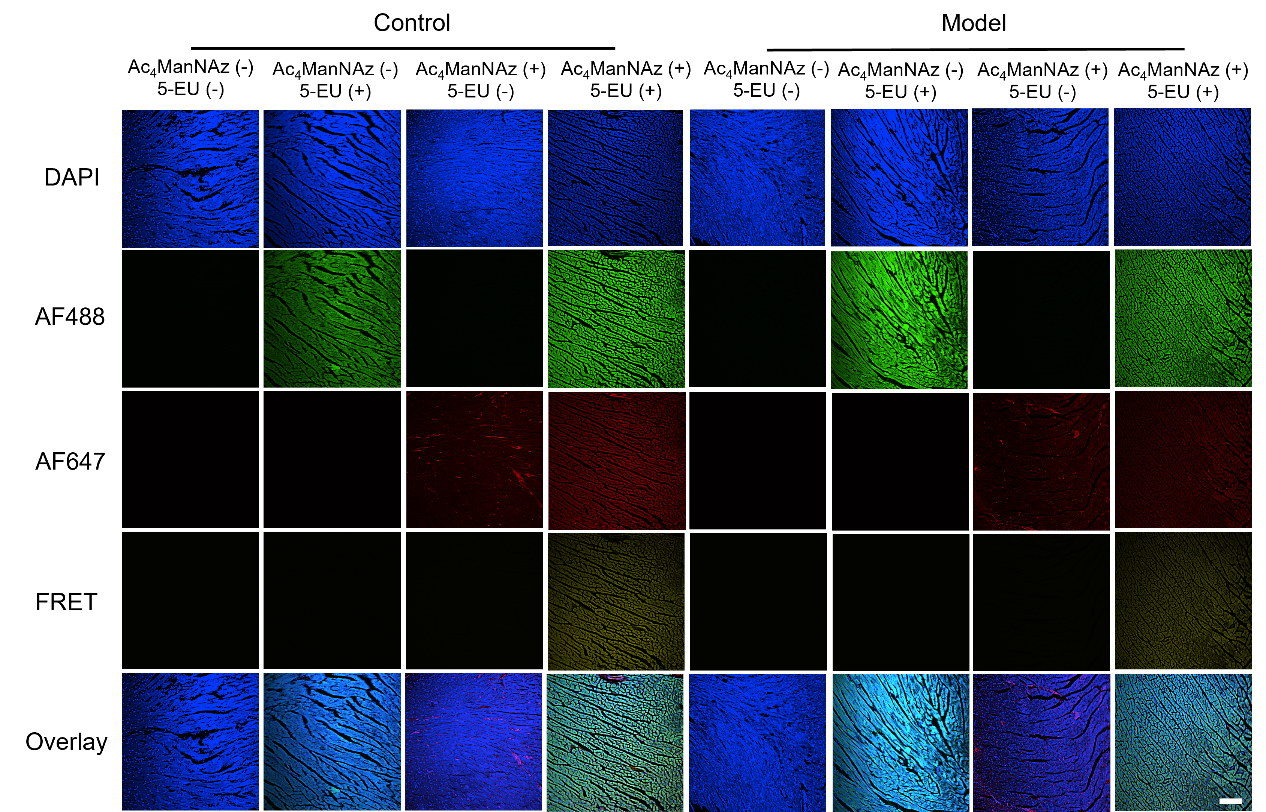

**Supplementary Fig. 33 |** **CLSM images of glycoRNAs in the heart section of different treated mice by COMPASS.**

The experimental mice were divided into the following two groups (n = 3 per group): a control group and the model group. To establish tumor metastasis models, 4T1 cells (1×10^7^ cells) suspended in PBS were injected intravenously through the tail vein in the model group. Meanwhile, the control group was injected with PBS. Mice were labeled with a single or two MCRs (100 μl of 20 mg/ml 5-EU, 0.16 mmole/kg DBCO-AF647), respectively, and subsequent slices were subjected to two-step click chemistry to achieve AF488 and AF647 labeling. Scale bar, 100 µm.

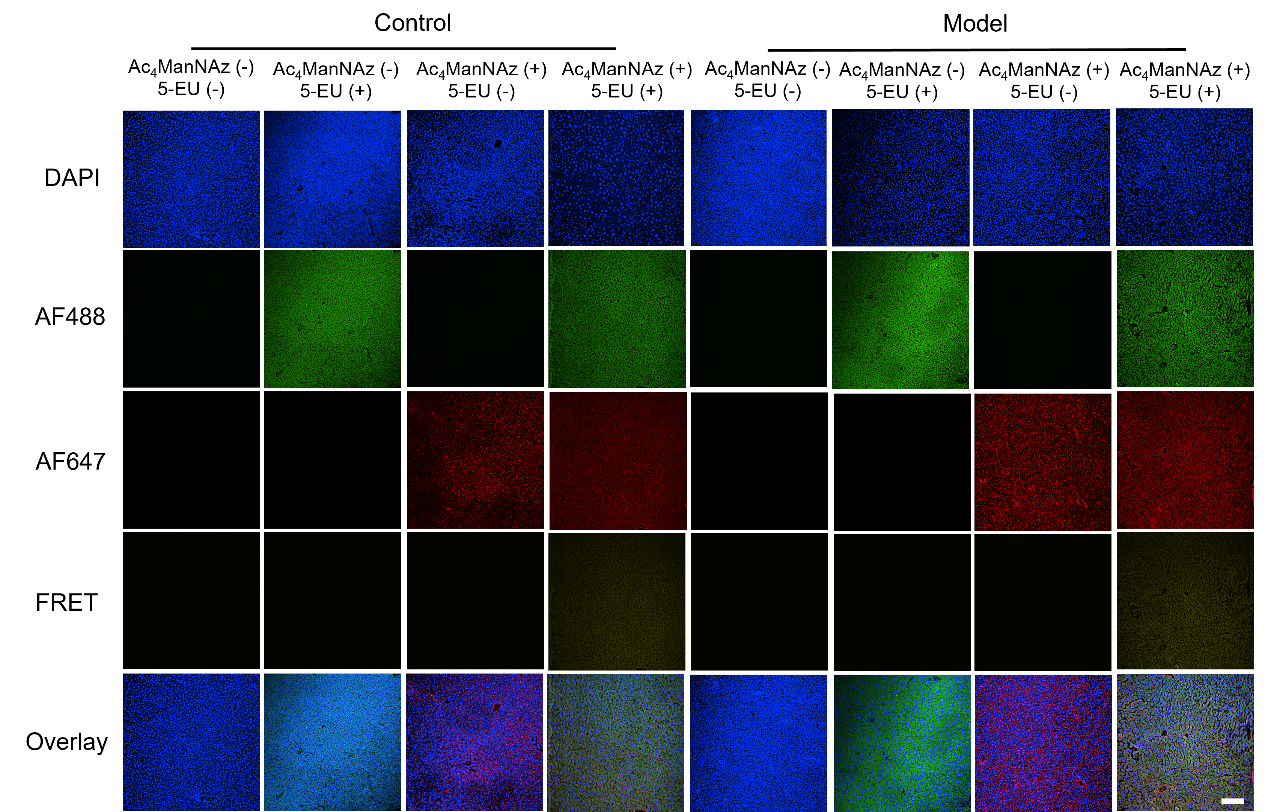

**Supplementary Fig. 34 |** **CLSM images of glycoRNAs in the liver section of different treated mice by COMPASS.**

The experimental mice were divided into the following two groups (n = 3 per group): a control group and the model group. To establish tumor metastasis models, 4T1 cells (1×10^7^ cells) suspended in PBS were injected intravenously through the tail vein in the model group. Meanwhile, the control group was injected with PBS. Mice were labeled with a single or two MCRs (100 μl of 20 mg/ml 5-EU, 0.16 mmole/kg DBCO-AF647), respectively, and subsequent slices were subjected to two-step click chemistry to achieve AF488 and AF647 labeling. Scale bar, 100 µm.

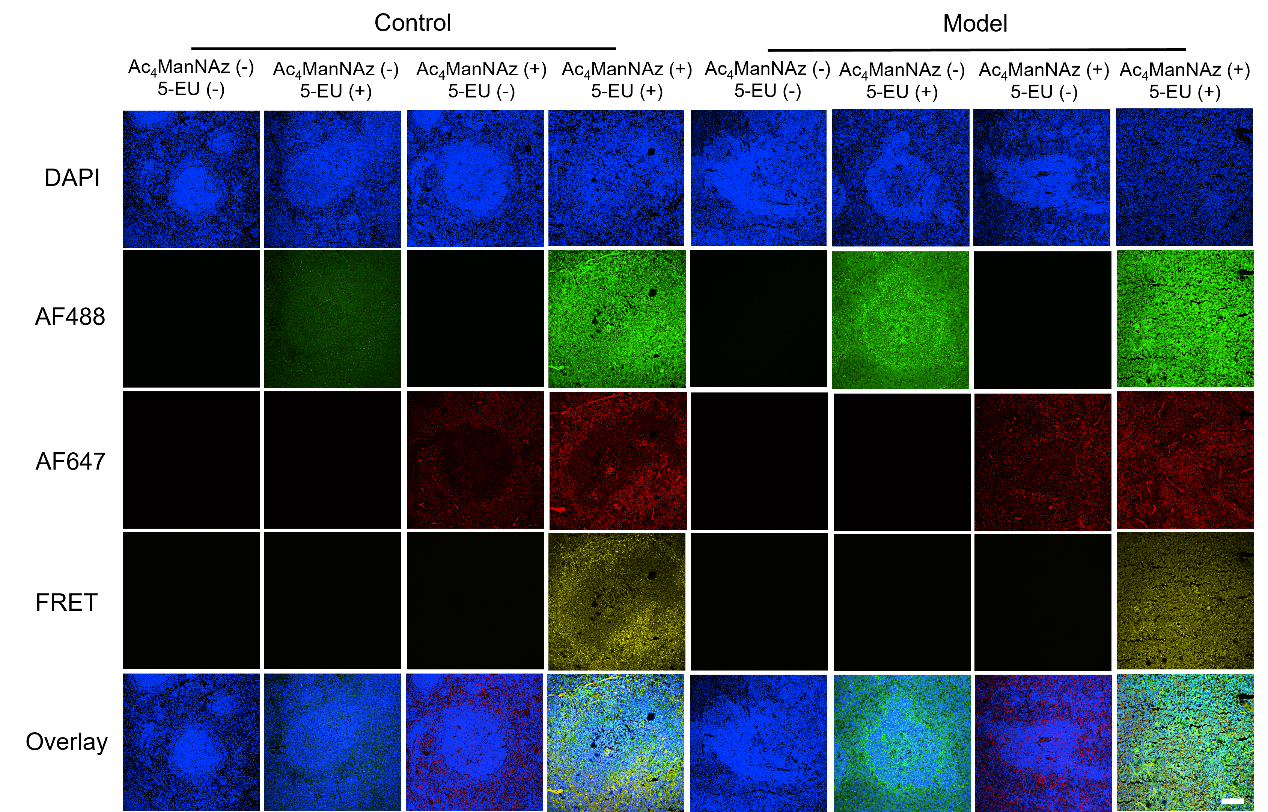

**Supplementary Fig. 35 |** **CLSM images of glycoRNAs in the spleen section of different treated mice by COMPASS.**

The experimental mice were divided into the following two groups (n = 3 per group): a control group and the model group. To establish tumor metastasis models, 4T1 cells (1×10^7^ cells) suspended in PBS were injected intravenously through the tail vein in the model group. Meanwhile, the control group was injected with PBS. Mice were labeled with a single or two MCRs (100 μl of 20 mg/ml 5-EU, 0.16 mmole/kg DBCO-AF647), respectively, and subsequent slices were subjected to two-step click chemistry to achieve AF488 and AF647 labeling. Scale bar, 100 µm.

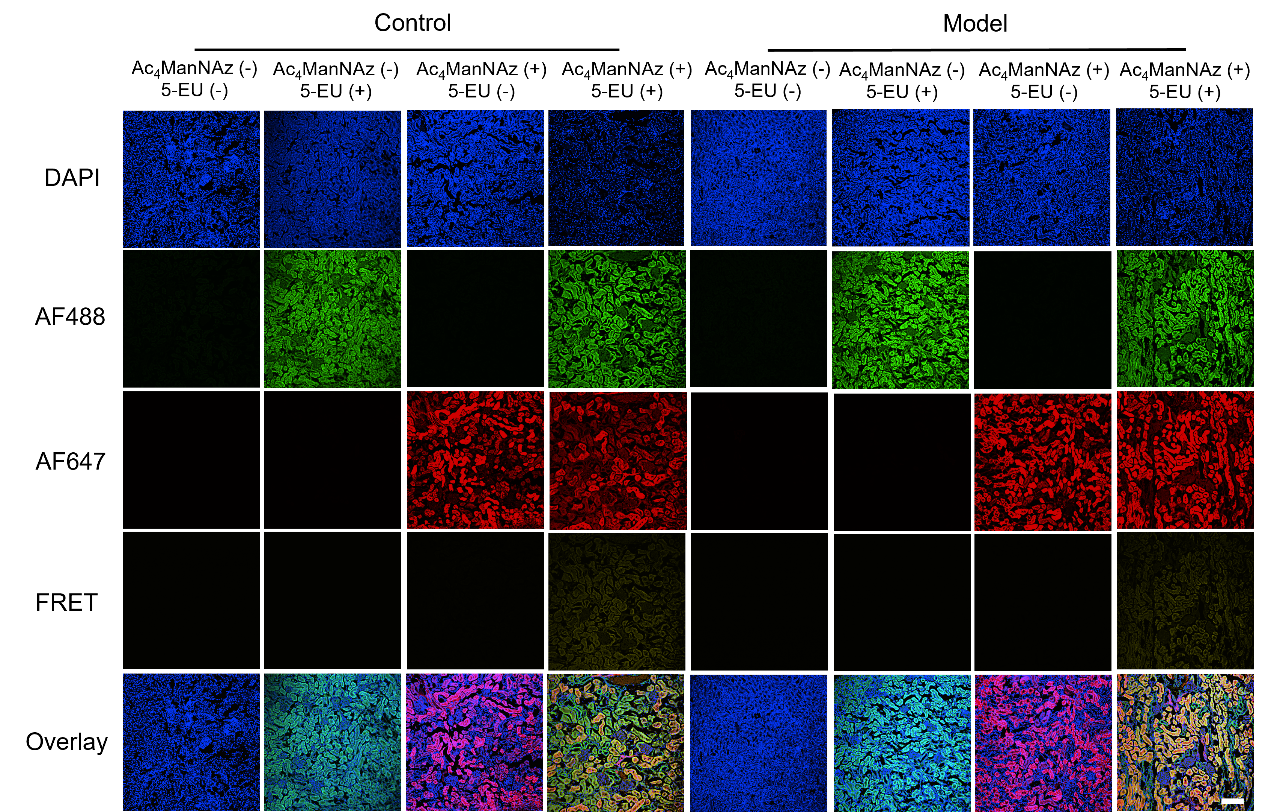

**Supplementary Fig. 36 |** **CLSM images of glycoRNAs in the kidney section of different treated mice by COMPASS.**

The experimental mice were divided into the following two groups (n = 3 per group): a control group and the model group. To establish tumor metastasis models, 4T1 cells (1×10^7^ cells) suspended in PBS were injected intravenously through the tail vein in the model group. Meanwhile, the control group was injected with PBS. Mice were labeled with a single or two MCRs (100 μl of 20 mg/ml 5-EU, 0.16 mmole/kg DBCO-AF647), respectively, and subsequent slices were subjected to two-step click chemistry to achieve AF488 and AF647 labeling. Scale bar, 100 µm.

**Supplementary Table 1.** The comparison between COMPASS and existing glycoRNA imaging methods

| **Methods** | **ARPLA**^3^ | **HieCo-2**^4^ | **COMPASS (This work)** |
| --- | --- | --- | --- |
| **Strategy for RNA recognition** | Complementary probes | Complementary probes | **Metabolic reporters** |
| **Targeted GlycoRNA Types** | U1, U35a, U8, and Y5 glycoRNAs  (Identified sequences) | U1, U3, and Y5 glycoRNAs  (Identified sequences) | **GlycoRNAs with U**  **(Compatible with unknown sequences)** |
| **Reaction Types** | Hybridization, ligation, DNA polymerization  (Environmental-sensitive, slow) | Hybridization  (Environmental-sensitive, slow) | **Click chemistry**  **(Robust, rapid)** |
| **Reagent Accessibility** | Customized DNA probes, multiple enzymes  (Specialized skills for design and operate) | Customized DNA probes  (Specialized skills for design and operate) | **Commercialized chemicals**  **(Stable, high accessibility)** |
| **Timing (from cell culture completion to imaging)** | ~ 8 h | ~ 4.5 h | **~ 1 h** |
| **Fluorogenic Component** | FAM  (λ_ex_ = 488 nm, λ_em_ = 525 nm, autofluorescence-sensitive) | AF647  (λ_ex_ = 650 nm, λ_em_ = 671 nm, autofluorescence-sensitive) | **FRET**  **(λ_ex_ = 488 nm, λ_em_ = 671 nm, autofluorescence-resistant)** |
| **GlycoRNA Signal Origination** | Signals from long DNA chains generated by RCA | Signals from long DNA chains generated by HCR | **Signals from dyes conjugated to glycoRNAs via short linkers** |
| **Sample Compatibility** | Cells | Cells | **Cells, organoids, tissues**  **(First visual evidence of glycoRNA distribution in tissues)** |
